# Innate Immune Dysregulation Precedes Clinical Onset in Neuromyelitis Optica

**DOI:** 10.64898/2026.10.16.757427

**Authors:** Leonie Müller-Jensen, Debanjana Chakravarty, Gianmarco Bellucci, Yuan Ding, Michelle Hughes, Alejandro M. Garcia, Carson E. Moseley, Ravi Dandekar, John W. Rose, Lisa K. Peterson, Ka-Ho Wong, Bradley Demarest, Noel Carlson, David E. Horvat, Keith Groshans, Nathan Tagg, Spencer R. Osbourn, Sithembile L. Mabila, Ruba Bou-Chahine, Victor Wang, M. Mateo Paz Soldan, Sean J. Pittock, Bruce A. C. Cree, Scott S. Zamvil, Ari J. Green, Adil Harroud, Sergio E. Baranzini, Michael R. Wilson, Stacey L. Clardy, Ahmed Abdelhak, Tammy L. Smith

## Abstract

Neuromyelitis optica spectrum disorder (NMOSD) is a disabling inflammatory central nervous system (CNS) disease characterized by pathogenic aquaporin 4 (AQP4) autoantibodies. The biological events that precede clinical onset remain unknown. In a unique longitudinal NMOSD cohort from the U.S. Department of Defense Serum Repository, with samples collected up to 24 years before onset, we applied discovery proteomics (∼5,400 proteins), pan-viral antibody profiling (VirScan), and *in silico* analyses to characterize preclinical NMOSD versus matched healthy controls. Innate immune activation, particularly CXCL1 elevation, was the dominant immunological feature of preclinical NMOSD, preceding onset by ∼12.7 years. AQP4-IgG was detectable more than 3 years before onset, while CNS injury markers increased only after disease manifestation. VirScan identified enhanced CMV-specific antibody reactivity following AQP4-IgG seroconversion, and network analyses revealed interactions between NMOSD-associated and CMV-derived proteins. Together, these findings establish a novel multi-hit model of NMOSD, with implications for understanding antibody-mediated autoimmunity more broadly.

## Introduction

Neuromyelitis optica spectrum disorder (NMOSD) is a relapsing, disabling, and sometimes fatal autoimmune disease of the central nervous system (CNS).^1,2^ Central to its pathogenesis are autoantibodies directed against the water channel aquaporin-4 (AQP4), which initiate AQP4 degradation, antibody-dependent cellular cytotoxicity, and complement activation,^3,4^ resulting in devastating inflammatory lesions in the optic nerves, spinal cord, and brain.^5^ AQP4 autoantibodies (AQP4-IgG) primarily originate from autoreactive plasmablasts in the periphery,^6,7^ and disease manifestation is therefore thought to depend on their access to the CNS, where they bind AQP4 on astrocyte end-feet.^8^

Despite advances in understanding NMOSD pathophysiology, current concepts rely largely on animal models that capture only selected aspects of disease and human samples collected after clinical onset.^9,10^ As such, the biological events that initiate the pathogenic cascade have remained unclear. It is unknown whether NMOSD is preceded by a prodromal phase of subclinical neuronal injury, as recently established for multiple sclerosis (MS),^11,12^ and whether infectious triggers may contribute to disease initiation. Moreover, NMOSD often co-occurs with other autoimmune diseases such as systemic lupus erythematosus (SLE) and Sjögren’s disease (SjD),^13^ but how immune perturbations in these conditions relate to NMOSD remains poorly understood.

Here, we leveraged the U.S. Department of Defense Serum Repository (DoDSR) to identify a unique longitudinal cohort of individuals who, in the years following sample collection, developed NMOSD. Timing of AQP4-IgG seroconversion was determined using serum samples collected up to 24 years before disease onset. Proximity extension proteomics quantifying ∼5,400 proteins was performed to identify differentially expressed proteins (DEPs) in preclinical NMOSD relative to matched healthy controls (HC). Potential microbial triggers were assessed using Virscan, a pan-viral antibody profiling method based on phage immunoprecipitation sequencing (PhIP-Seq), and network-based *in silico* approaches.

Together, this work defines a prolonged preclinical phase of innate immune activation in NMOSD that precedes AQP4-IgG seroconversion, CNS injury, and clinical onset by multiple years. We implicate cytomegalovirus (CMV) as a potential contributor to disease development and identify a serum proteome signature associated with NMOSD, collectively providing new insight into the earliest stages of antibody-mediated autoimmunity and suggesting a potential window for earlier diagnosis and treatment.

## Results

### Cohort description

In total, 267 samples from 33 AQP4-IgG+ NMOSD cases (70% female, 33% ≥ 40 years old) and 57 HC (63% female, 35% ≥ 40 yrs old) were included in the primary analysis (**Figure 1**). Demographics and cohort characteristics are summarized in **Table 1**. AQP4-IgG autoantibodies were first detected a median of 3.4 years (IQR, 8.7 to 1.3 years) before clinical onset (**Figure 2a**). In 11 of 33 patients (33.3%) the first available serum sample, collected at a median of 12.7 years (IQR, 14.4 to 6.6 years) before onset, was already positive for AQP4-IgG autoantibodies, indicating that the time between AQP4-IgG seroconversion and clinical onset was even longer than our estimate. Closer to disease onset, an increase in AQP4-IgG titres was observed (**Figure 2b, Supplemental Figure 1**).

**Figure 1.**
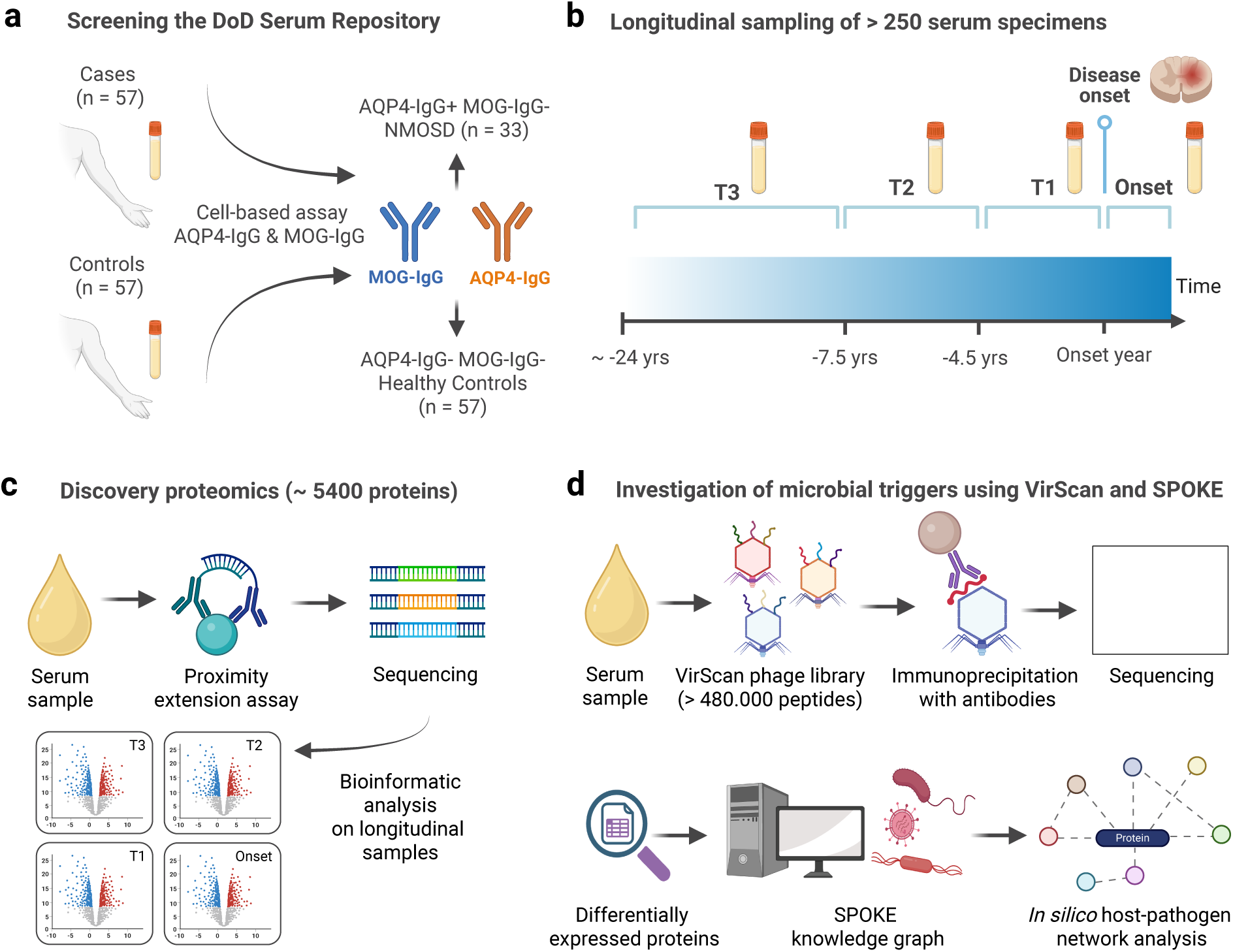
Study overview. **(a)** Schematic of Department of Defense (DoD) Serum Repository study cohort. 57 NMOSD cases and 57 age- and sex-matched healthy controls were identified. As diagnostic antibody assays were not specified, AQP4-IgG and MOG-IgG were retrospectively assessed by fixed cell-based assay. This confirmed 33 cases of AQP4-IgG+ NMOSD; the remaining 24 cases were either autoantibody negative or MOG-IgG+ and were excluded from the primary analysis. **(b)** Overview of the four investigated time intervals relative to disease onset (T1/T2/T3, Onset). **(c, d)** *In vitro* and *in silico* analyses performed on longitudinal serum samples. SPOKE, Scalable Precision Medicine Open Knowledge Engine.

**Table 1.** Demographics and clinical characteristics of the DoDSR cohort.

| Variable | Healthy Controls<br>(n=57) | NMOSD<br>(n=33) | P-value |
| --- | --- | --- | --- |
| **Biological sex, n (%)** |  |  | 0.647 |
| Female | 36 (63) | 23 (70) |  |
| Male | 21 (37) | 10 (30) |  |
| **Age at onset, n (%)** |  |  | 1.000 |
| < 40 years | 37 (65) | 22 (67) |  |
| ≥ 40 years | 20 (35) | 11 (33) |  |
| **Total no. of samples** | 165 | 102 |  |
| Pre-onset | 133<sup>a</sup> | 82 |  |
| Post-onset | 32<sup>a</sup> | 20 |  |
| **Presenting symptom location, n (%)<sup>b</sup>** |  |  |  |
| Spinal cord | - | 19 (58) |  |
| Optic nerve | - | 12 (36) |  |
| Brain (supra- and infratentorial) | - | 4 (12) |  |
| Area postrema | - | 4 (12) |  |
| Unknown | - | 1 (3) |  |
| **Concurrent autoimmune disease n (%)<sup>c</sup>** |  |  |  |
| SjD or known anti-SSA autoantibodies | - | 6/24 (25) |  |
| SLE | - | 2/24 (8) |  |
| Hashimoto's disease | - | 1/24 (4) |  |
| Other<sup>d</sup> | - | 1/24 (4) |  |
| **Disease-modifying therapy after onset<sup>#</sup>** |  |  |  |
| **RTX** | - | 20/32 (63) |  |
| **AZT** | - | 13/32 (41) |  |
| **MMF** | - | 6/32 (19) |  |
| **MTX** | - | 2/32 (6) |  |
| **CTX** | - | 1/32 (3) |  |

**Figure 2:**
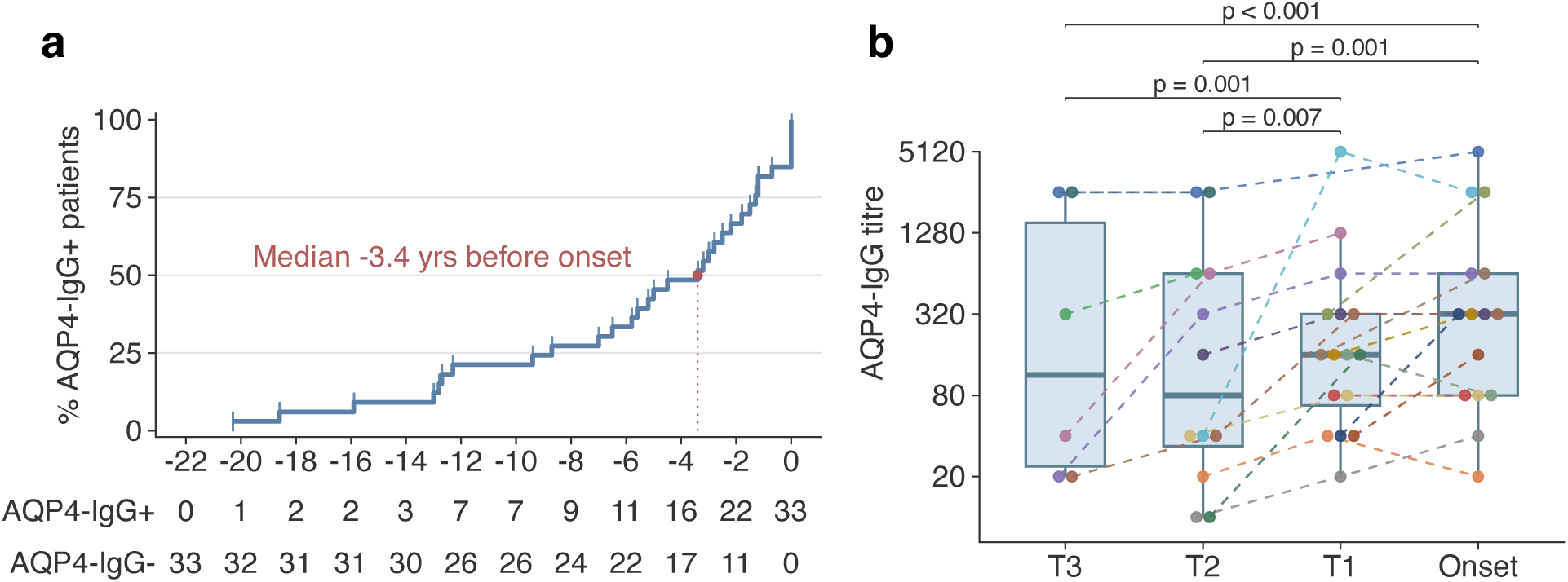
Timing of first AQP4-IgG autoantibody detection and titre trajectories. **(a)** Empirical Cumulative Distribution Function (ECDF) showing timing of first AQP4-IgG+ serum sample relative to disease onset. X-axis denotes time in years (yrs) before clinical onset, y-axis shows the cumulative percentage of patients who had seroconverted. For 5 patients, the first sample with detectable AQP4-IgG autoantibodies were the post-onset samples. These patients were assigned x = 0 for visualization purposes. **(b)** AQP4-IgG titres stratified by time intervals relative to disease onset. Samples collected >7.5 years before onset were classified as T3, those collected 7.5-4.5 years before onset as T2, those collected within 4.5 years before onset as T1, and those collected within 90 days of first presentation as onset. Boxplots show the median, interquartile range (IQR; box), and 1.5 × IQR (whiskers). Colors indicate individual patients. Patient-specific titre trajectories are connected by dashed lines. P-values are derived from a linear mixed model restricted to samples that were AQP4-IgG+ and where > 1 sample per patient was available (n = 19 patients, n = 49 samples).

### Systemic immune dysregulation precedes clinical onset in AQP4-IgG+ NMOSD

Longitudinal comparison of all available samples from NMOSD cases (pre- and post-onset) and HC identified 121 DEPs (false discovery rate [FDR] 5%) (**Figure 3a, Supplemental Table 1**). Out of the top five proteins, CXCL1 (log_2_FC 1.16, p_adj_ < 0.001), CXCL3 (log_2_FC 1.32, p_adj_ < 0.001) and CXCL5 (log_2_FC 0.97, p_adj_ < 0.001) - all ligands of the neutrophil receptor CXCR2 - as well as the T cell chemoattractant CXCL11 (log_2_FC 0.90, p_adj_ < 0.001) were increased in NMOSD versus HC, whereas CCL24 was reduced (log_2_FC -0.74, p_adj_ < 0.001).

**Figure 3:**
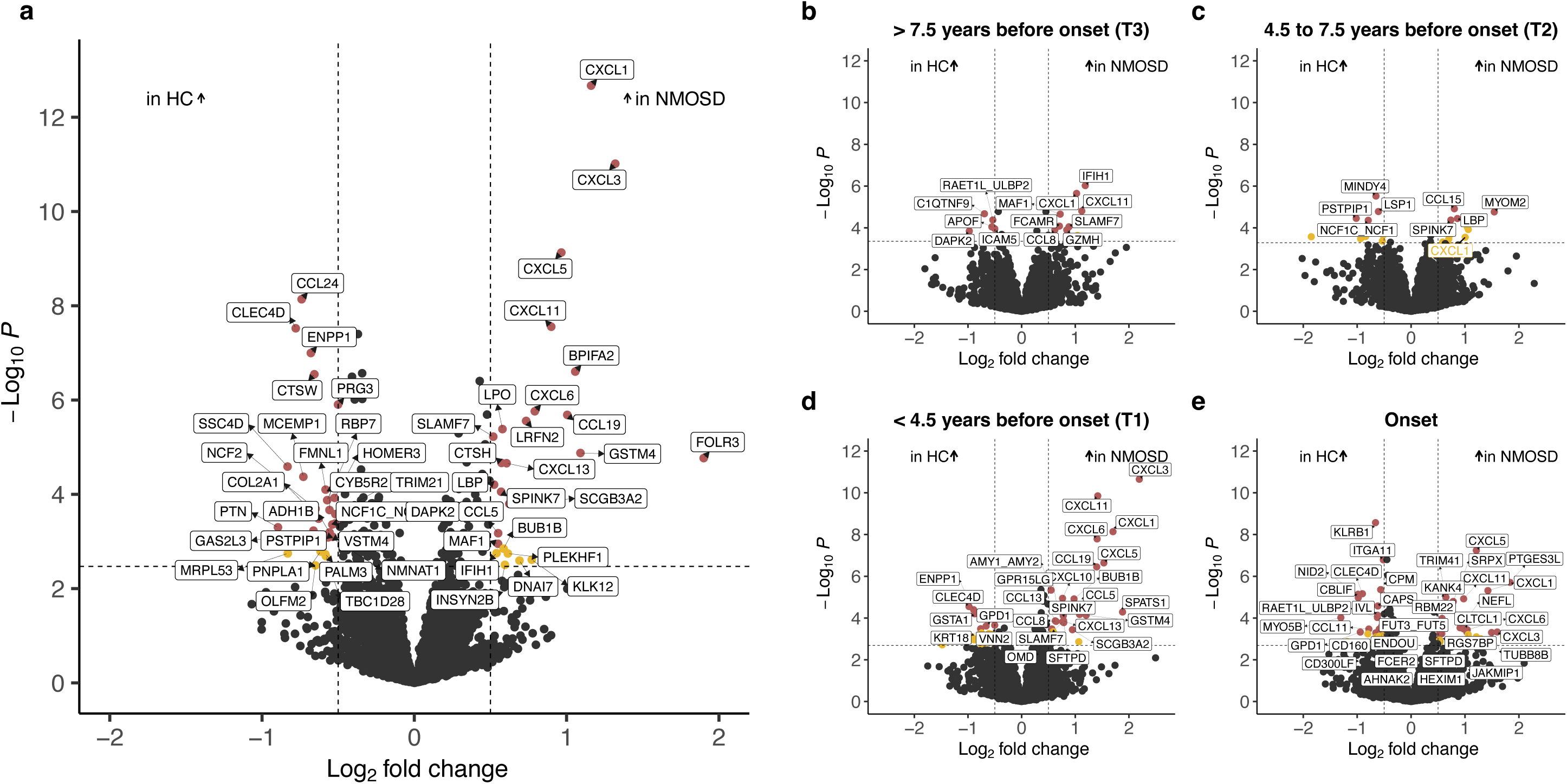
Differentially expressed serum proteins (DEPs) between patients with NMOSD and HC. **(a)** Volcano plot showing DEPs between NMOSD cases and HC combining pre- and post-onset samples (NMOSD: n = 102, HC: n = 165). **(b-e)** DEPs at different time intervals relative to disease onset. For duplicate samples within a time interval and patient (n = 17 HC and n = 9 NMOSD cases), samples were merged into a single mean value per assay. **(b)** Included n = 36 HC and n = 24 NMOSD samples, **(c)** n = 38 HC and n = 24 NMOSD samples, CXCL1 is highlighted in yellow (p_adj_ = 0.073, log_2_FC = 1.00), **(d)** n = 42 HC and n = 25 NMOSD samples, **(e)** n = 32 HC and n = 20 NMOSD samples. DEPs at FDR 5% and 10% with a log_2_FC > 0.5 are highlighted in red and yellow, respectively. Labels are restricted to significant DEPs at FDR 5% and log_2_FC > 0.5 in panel **b-e**, except for CXCL1 in **c**. Vertical and horizontal dashed lines indicate a threshold of log_2_FC > 0.5 and FDR 10%, respectively.

To characterize proteome changes preceding the clinical onset of NMOSD, DEPs across distinct time intervals were analyzed (**Figure 3b-e**). Since up to four consecutive samples were available per patient (median 3, range 2-4), we defined four intervals, maximizing the number of patients contributing one sample per interval (**Supplemental Figure 2**): (1) samples collected within 90 days of disease onset or after (Onset), (2) samples collected ≤ 4.5 years before clinical onset (T1), (3) samples collected > 4.5 but ≤ 7.5 years before clinical onset (T2), and (4) samples collected earlier than 7.5 years before clinical onset (T3). Comparing the earliest samples (T3; median, 12.7 yrs before onset [IQR, 17.9 to 9.4]) with time-matched HC, we identified 16 DEPs (FDR 5%). Again, CXCL1 (log_2_FC 1.02, p_adj_ = 0.006) and CXCL11 (log_2_FC 1.12, p_adj_ = 0.017) ranked among the top five proteins. Although the number of DEPs transiently decreased to 8 at T2, the extent of proteome changes increased progressively as NMOSD onset approached, rising to 46 DEPs at T1 and 60 DEPs at clinical onset. At FDR 10%, CXCL1 was the only protein that showed persistent elevation across all timepoints evaluated (**Figure 3b-e**). At FDR 25%, this was also observed for CXCL3, CD207, and ICAM5 **(Supplemental Figure 3a-d).**

Given the variability in the timing of the first autoantibody-positive serum sample, ranging from 20 years before disease onset to after clinical manifestation, proteome changes were examined specifically during the pre- and post-seroconversion period of the preclinical stage, independent of their time interval. Notably, CXCL1, CXCL3, and CXCL5 were consistently elevated in NMOSD compared to HC, independent of AQP4-IgG seroconversion (**Supplemental Figure 4a-b**). By contrast, CCL19 - a chemokine promoting lymphocyte migration to the thymus and secondary lymphoid organs^14^ - emerged only post-seroconversion and was then the most prominent DEP (log_2_FC 1.67, p_adj_ < 0.001). As an additional finding, lower levels of TRIM21, a known antigen in SjD and SLE,^15^ were observed following AQP4-IgG seroconversion (log_2_FC - 0.25, p_adj_ = 0.062). Seven patients with a preexisting diagnosis of SjD, SLE, or detectable anti-SSA autoantibodies showed particularly low TRIM21 levels, which decreased over time (**Supplemental Figure 5a-b)**.

### Elevation of CNS injury markers is restricted to clinical onset in NMOSD

Next, in a hypothesis-driven approach, we examined trajectories of markers associated with astrocyte damage (glial fibrillary acidic protein [GFAP]), neuroaxonal injury (neurofilament light chain [NEFL]), and myelin injury (myelin oligodendrocyte glycoprotein [MOG]). Both GFAP and NEFL were increased following clinical onset, with higher GFAP (log_2_FC = 0.69, p_nom_ = 0.007) and NEFL levels (log_2_FC = 0.91, p_nom_ < 0.001) in NMOSD cases versus HC. Moreover, higher AQP4-IgG titres post-onset were associated with higher GFAP levels (log_2_FC 1.04, p_nom_ = 0.003), while NEFL showed no significant association (log_2_FC 0.22, p_nom_ = 0.056). By contrast, serum NEFL and GFAP levels did not differ between NMOSD cases and HC at any preclinical interval (NEFL: T1: log_2_FC -0.02, p_nom_ = 0.892; T2: log_2_FC 0.17, p_nom_ = 0.352; T3: log_2_FC -0.04, p_nom_ = 0.820; GFAP: T1: log_2_FC 0.16, p_nom_ = 0.267; T2: log_2_FC 0.31, p_nom_ = 0.059; T3: log_2_FC 0.04, p_nom_ = 0.777; **Supplemental Figure 6a-b**). Differences in MOG protein abundance were not detected at any timepoint.

### Limited evidence of B cell and complement activation during preclinical NMOSD

As autoantibody-mediated complement activation and T helper 17 cell (Th17) responses are central features of NMOSD immunopathology,^16,17^ proteins related to B-cell homeostasis (B-cell maturation antigen [BCMA], B-cell activating factor [BAFF], B-cell activating factor receptor [BAFF-R], CXCL13), complement (C1r/s, C3, C4b, C5, C7, C8b, C9), and Th17 differentiation (IL6, IL17A, IL23R) were evaluated. Consistent with B-cell recruitment and plasma cell expansion, CXCL13 and BCMA showed modest but gradually increasing elevations throughout the preclinical period (**Supplemental Figure 7a-b**). BAFF exhibited a transient increase at T1 (log_2_FC = 0.15, p_nom_ = 0.041), while serum abundance of the soluble BAFF-R was comparable between preclinical NMOSD and HC (**Supplemental Figure 7c-d**). Among complement components, C7 was the only protein that showed a significant increase at T1 (log_2_FC = 0.36, p_nom_ = < 0.001) and remained elevated thereafter (log_2_FC = 0.34, p_nom_ = 0.003; **Supplemental Figure 7e**). With regard to Th17 signaling, only IL23R was reduced in preclinical NMOSD cases compared with HC at T1 (log_2_FC = -0.35, p_nom_ = 0.037), whereas IL6 and IL17A showed no differences between the groups at any time interval (**Supplemental Figure 8**).

### Early and sustained enrichment of innate immune pathways in NMOSD

To place the observed immune signatures into a broader biological context, we performed pathway enrichment analyses by mapping identified DEPs to curated pathways from the Reactome and Gene Ontology Biological Process (GO:BP) databases.^18,19^

Across all preclinical timepoints, pathways related to chemokine signaling, antimicrobial humoral immune responses, and granulocyte migration were strongly upregulated **(Supplemental Figure 9a-b)**. Within the adaptive immune response, pathways involved in T-cell activation and migration were more frequently enriched than pathways related to B-cell responses. During the preclinical stages, T-cell-related pathways accounted for 3/19 (T3), 18/210 (T2), and 7/147 (T1) of significantly upregulated GO:BP terms. In contrast, B cell-related pathways were absent in the earliest time window (T3) and limited to two enriched pathways (“*B cell activation*” and “*B cell-mediated immunity*”) emerging closer to disease onset.

### Genetic variants in the *CXCL1* gene are not associated with NMOSD

Having identified signs of systemic immune dysregulation in patients who later developed NMOSD, we next explored potential drivers of this process. First, given its consistent enrichment in NMOSD across all time intervals, we investigated whether elevated CXCL1 reflected genetic susceptibility to NMOSD. To that end, we tested the *CXCL1* locus using gene-based analysis (MAGMA).^20^ Genome-wide association study (GWAS) was performed in the UK Biobank (UKB) and All of Us (AoU) cohorts and meta-analyzed with two previously published NMOSD GWAS^21,22^ (**Supplemental Table 2**). In a total of 538 cases, no association was found (p_nom_ = 0.538). Inclusion of another recently published cohort^23^ increased the sample size to 2,111 NMOSD cases of diverse ancestry and did not change the result (p_nom_ = 0.750). Collectively, these findings provide no genetic support for a role of *CXCL1* polymorphisms in NMOSD susceptibility, indicating that its early elevation is unlikely to be driven by common genetic variation.

### Enhanced antibody responses against CMV emerge after AQP4-IgG seroconversion

In MS, increasing evidence suggests that Epstein-Barr virus (EBV) infection is a prerequisite for disease initiation.^24^ Hypothesizing that specific microorganisms may similarly be associated with NMOSD pathogenesis, we examined the relationship between AQP4-IgG seroconversion and antiviral reactivity patterns. We performed VirScan,^25^ a PhIP-Seq technology for profiling antibody responses against > 480,000 viral peptides. A paired analysis of samples collected pre- and post-AQP4-IgG seroconversion (n = 20 each) identified 7 viral peptides with differential reactivity (FDR 5%) that were also elevated relative to time-matched HC (**Figure 4a**). Whereas 2 peptides were enriched pre-seroconversion, 5 exhibited increased reactivity post-seroconversion, all of which were derived from herpesviruses, specifically CMV (also named human herpesvirus 5 [HHV5]) and human herpesvirus 2 (HHV2) (**Figure 4b**).

**Figure 4.**
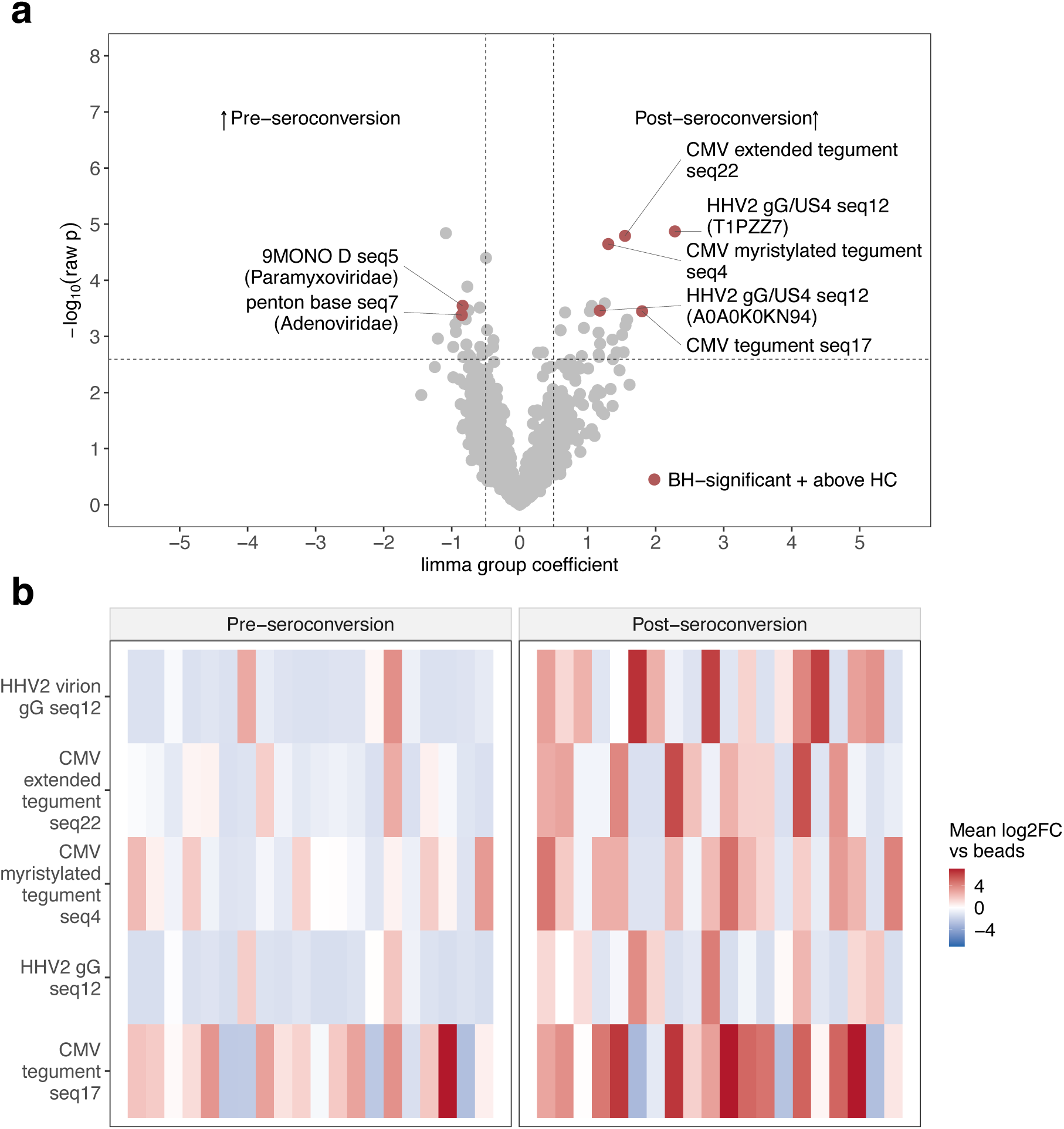
Comparison of pre- and post-seroconversion antiviral reactivity patterns in NMOSD. **(a)** Volcano plot showing peptide-level antiviral antibody reactivity in 20 NMOSD cases with paired pre- and post-seroconversion samples. Differential reactivity was tested using a paired *limma* model applied to sample-level mean log_2_FC relative to beads. Red dots indicate peptides that were significant at FDR 5% and demonstrated higher signal than time-matched HC. Vertical and horizontal dashed lines mark an absolute *limma* coefficient of 0.5 and the nominal significance threshold, respectively. **(b)** Heatmap of the five peptides enriched after AQP4-IgG seroconversion. Each row represents one peptide, and each column represents one patient sample. Color indicates mean log_2_FC versus beads.

Consistent with an enhanced antibody response against CMV, NMOSD cases demonstrated a greater increase in overall anti-CMV reactivity across the AQP4-IgG seroconversion period compared to a longitudinal analysis of HC samples. Specifically, the CMV reactivity burden - defined as the number of CMV peptides enriched above beads - exhibited a marked increase following AQP4-IgG seroconversions in NMOSD but remained largely unchanged over time in HC (4.9 versus 0.8 peptides, p_nom_ interaction = 0.008). Similarly, the normalized CMV peptide signal demonstrated a higher enrichment following AQP4-IgG seroconversion in NMOSD compared to longitudinal changes in HC (increase in log_10_[1 + normalized mean reads per 100,000] of 0.31 versus 0.11, p_nom_ interaction = 0.048). Thus, the period of AQP4-IgG seroconversion was associated with a marked increase in anti-CMV reactivity that exceeded longitudinal fluctuations observed in HC. In contrast, this differential longitudinal response between NMOSD and HC was not observed for EBV, since overall anti-EBV reactivity increased in both groups (**Supplemental Figure 10a-d**). Post-onset comparisons between NMOSD patients and HC did not identify viral peptides with FDR-significant differential reactivity.

### *In silico* network analysis links proteome changes in preclinical NMOSD to CMV

Infections influence the human proteome.^26^ To investigate potential interactions between the identified proteome changes in preclinical NMOSD and specific microorganisms, we performed an *in silico* analysis using the Scalable Precision Medicine Open Knowledge Engine (SPOKE) knowledge graph, with DEPs serving as seeds.

Again, a CMV network signature emerged as the most prominent viral signal across different disease stages (**Supplemental Figure 11a-c**). Both pre-seroconversion (z = 13.05, p_adj_ < 0.001) and post-onset (z = 7.81, p_adj_ < 0.001) CMV exhibited the highest proximity score among all nodes. During the post-seroconversion period, CMV remained the top-ranking viral signal, although it did not reach statistical significance (z = 3.00, p_adj_ = 0.28).

Beyond CMV, a consistent bacteria-biased network signature was observed throughout the NMOSD disease trajectory (**Supplemental Figure 11d),** with several bacterial species (spp.) showing persistent enrichment across timepoints. Specifically, *Porphyromonas gingivalis* was detected as a significant node during the preclinical stages (z = 3.28; p_adj_ = 0.047 and z = 3.71; FDR = 0.035, for pre- and post-seroconversion, respectively), while *Listeria monocytogenes* emerged after AQP4-IgG4 seroconversion (z = 4.83; p_adj_ < 0.001 and z = 7.04; FDR < 0.001 for post-seroconversion and post-onset, respectively). After disease onset, an even broader bacterial signal emerged. In addition to *L. monocytogenes*, mucosa-associated pathogens ranked among the top hits, including *Streptococcus pyogenes* (z = 5.13) and *Clostridium spp. (*z = 5.05) **(Supplemental Figure 11a-c)**. Analysis of metapaths between microorganisms and NMOSD-associated DEPs revealed that human protein targets of *L. monocytogenes*, *Clostridium spp*., and CMV consistently converged on CXCL1 and CXCL3, with CMV showing the strongest connectivity (**Supplemental Figure 12a-c**).

### A 39-protein panel distinguishes preclinical NMOSD from HC

Given substantial proteome changes in individuals who subsequently developed NMOSD, we tested whether these signatures could distinguish NMOSD from HC using machine learning approaches. Boruta feature selection,^27,28^ applied to samples collected at or after disease onset, identified 39 proteins that discriminated NMOSD from HC **(Figure 5a).** We then trained multiple machine learning classifiers using this protein panel. Ridge regression with 5-fold cross-validation showed the best performance, achieving a mean AUC of 0.95 over 100 repeats (95% CI, 0.92-0.96) (**Figure 5b**). Model performance exceeded chance (permutation p < 0.001), and adding GFAP - a recognized biomarker of astrocytic injury in NMOSD^29^ - to the panel did not further improve model performance (AUC 0.95; 95% CI, 0.92-0.96, **Supplemental Table 3**). Importantly, the classifier maintained robust discrimination between NMOSD cases and HC when applied to the corresponding preclinical samples, including those collected > 7.5 years before clinical onset (T3; AUC 0.90; 95% CI, 0.82-0.98, **Figure 5b**).

**Figure 5:**
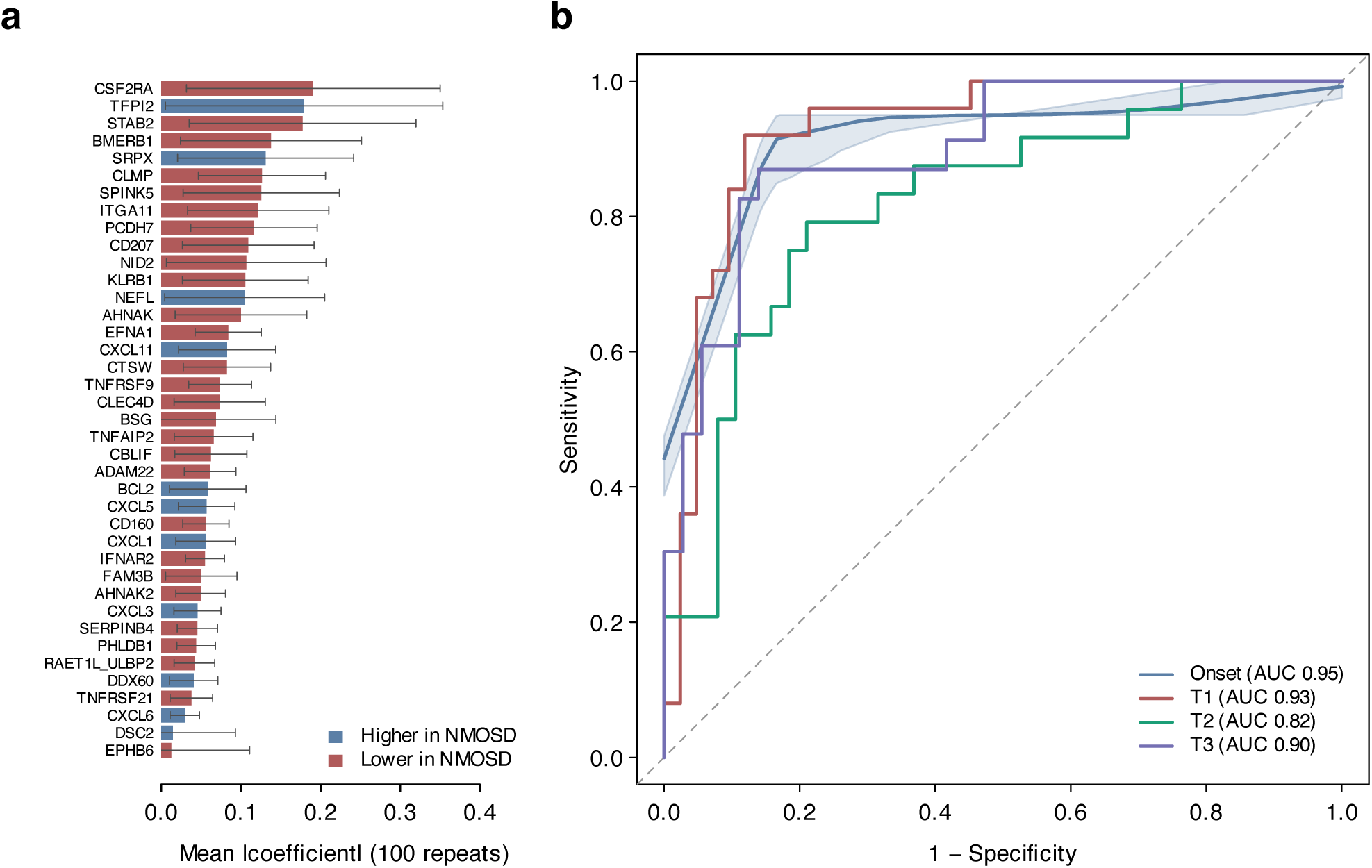
Machine learning classifier distinguishing preclinical NMOSD from HC. A penalized logistic regression model (Ridge regression) with 5-fold cross-validation was developed on post-onset samples from AQP4-IgG+ NMOSD cases (n = 20) and time-matched HC (n = 32) and applied to pre-onset samples. **(a)** Importance plot of 39 proteins selected in ≥ 60% of iterations using repeated Boruta feature selection. Bars represent the mean absolute coefficient across 100 classifier model repeats; error bars indicate ± 1 SD**. (b)** Receiver operating characteristic (ROC) curves for the mean-coefficient classifier. ROC at disease onset (blue) is shown with its 95% bootstrap confidence band (light blue shading); temporal replication curves indicate preclinical timepoints T1, T2, and T3.

### Distinct proteome signatures differentiate AQP4-IgG+ NMOSD from suspected seronegative cases

Approximately 10-20% of patients with clinical syndromes consistent with NMOSD lack detectable AQP4-IgG or MOG-IgG autoantibodies and are currently classified as seronegative NMOSD, though this disease category remains controvercial.^30^ In these patients, disease pathogenesis is less clear and established NMOSD therapies are less effective.^31^

To compare the serum proteome of AQP4-IgG+ NMOSD to cases of suspected seronegative disease, we analyzed 11 additional patients from the DoDSR (**Supplemental Figure 13**). Of these, all had developed inflammation of the optic nerve, spinal cord, or both and were seronegative for MOG-IgG and AQP4-IgG by cell-based assay (**Supplemental Table 4**). When including all available pre- and post-onset samples from both groups, two DEPs were identified at FDR 5%: the neuronal protein PRSS53 (log_2_FC -0.53, p_adj_ 0.012) and the chemokine CCL24 (log_2_FC -0.82, p_adj_ 0.012) showed higher abundance in patients with suspected seronegative compared to AQP4-IgG+ NMOSD (**Supplemental Figure 14**). As CCL24 was also among the DEPs identified when comparing AQP4-IgG+ NMOSD and HC, we assessed additional overlapping DEPs at an FDR of 10%. Fourteen of 24 DEPs (58.3%) were shared across both comparisons, including seven proteins consistently increased (CXCL1, KLK12, FCRL5, CCL19, BPIFA2, CXCL3, DNAI7) and seven proteins consistently decreased (CCL24, NMNAT1, MEP1B, FMNL1, GHR, CPM, PIGR) in AQP4-IgG+ NMOSD relative to HC and suspected seronegative cases.

Finally, we applied the 39-protein classifier to post-onset samples from AQP4-IgG+ and suspected seronegative NMOSD cases and observed good discrimination between the two groups (AUC, 0.83; 95% CI, 0.66-1.00; **Supplemental Figure 15**). Collectively, these findings support a unique proteome signature of AQP4-IgG+ NMOSD, with a set of proteins distinguishing it from both HC and disease controls.

## Discussion

The discovery of AQP4-IgG established NMOSD as an autoantibody-mediated astrocytopathy in which autoreactive plasmablasts are considered the principal drivers of disease.^6, 32^ The present study expands this paradigm, demonstrating that innate immune dysregulation emerges multiple years before AQP4-IgG seroconversion and more than a decade before onset of clinical disease. Specifically, persistent elevation of the chemokine CXCL1 was identified as a hallmark of the preclinical phase of NMOSD. CXCL1 is produced by a broad range of cell types in response to inflammation and tissue damage.^33,34^ Together with related DEPs such as CXCL3 and CXCL5, it primarily signals through the CXCR2 receptor to promote neutrophil chemotaxis, degranulation, and formation of neutrophil extracellular traps (NETs). In the context of CNS inflammation, including AQP4-IgG-mediated autoimmunity,^10^ CXCL1 is produced by astrocytes, neurons, and endothelial cells to promote neutrophil recruitment across the blood-brain barrier (BBB).^35,36^

Although neutrophils are increasingly recognized as important mediators of CNS injury during NMOSD relapses, they have previously been regarded as amplifiers rather than initiators of disease. Active CNS lesions show prominent neutrophil infiltration, and granulocyte activation markers - including CXCL1 - are enriched in the CSF of NMOSD patients during acute attacks.^37–39^ Moreover, it was recently shown that patients with NMOSD exhibit increased bone marrow granulopoiesis with higher output of ISG15+ CXCR2+ neutrophils.^40^ These neutrophils expressed high levels of BAFF, a growth factor that supports the survival of autoreactive B cells, suggesting a mechanistic link between aberrant granulopoiesis and B-cell autoimmunity. In the present study, we build on these findings and show that granulocyte dysregulation begins substantially earlier than previously recognized. Enrichment of CXCL1, CXCL3, and CXCL5 preceded elevation of BAFF, AQP4-IgG detection, and complement activation, suggesting that neutrophil-associated inflammation occurs prior to - and potentially fosters - adaptive immune dysregulation in NMOSD. Importantly, early innate immune activation preceding autoantibody formation and clinical onset has also been observed in related systemic autoimmune diseases, including SLE and SjD.^41,42^

As we found no evidence of a genetic association between *CXCL1* polymorphisms and NMOSD, the cause of sustained CXCL1 elevation remains unknown. One hypothesis is that pathogens or environmental exposures contribute to the development of NMOSD. Indeed, VirScan identified increased reactivity to CMV across the AQP4-IgG seroconversion interval, beyond longitudinal changes observed in HC. This patterns is consistent with amplification of a preexisting CMV-specific antibody response and may reflect renewed CMV antigen exposure through viral reactivation, reinfection, or impaired viral control around the time of AQP4-IgG seroconversion.^43^ Using SPOKE, we further identified strong interactions between NMOSD-associated and CMV-derived proteins, with CXCL1 showing multiple connections to viral components. The convergent identification of CMV across *in vitro* and *in silico* approaches is noteworthy, since CMV can reside in granulocyte precursor cells, suppress apoptosis in neutrophils,^44^ and produce a viral CXCL1 homolog (vCXCL1) which binds to CXCR2.^45^ Notably, neutrophils from NMOSD patients have been found to show increased survival in response to toxic stimuli *in vitro*, potentially linking CMV to neutrophil dysregulation in NMOSD.^46^ Beyond CMV, SPOKE identified an association between *Clostridium spp.* and clinically manifest NMOSD, building on previous studies on T-cell cross-reactivity between *C. perfringens* and AQP4.^17,47^ Together with a recent report of a shared core peptide between AQP4 and T-cell epitopes derived from both CMV and *C. perfringens*,^48^ these findings raise the possibility that diverse microbial exposures, including CMV reactivation and bacterial infection, may contribute to NMOSD pathogenesis.

Another key finding of this study is that AQP4-IgG seroconversion precedes clinical onset by multiple years, despite the established pathogenicity of these autoantibodies. This temporal dissociation supports a multi-hit model of NMOSD in which systemic immune dysregulation (*first hit*) is followed by AQP4-IgG formation (*second hit*), and a subsequent unknown trigger - potentially BBB breakdown - that leads to onset of clinical symptoms (*third hit*). Similar concepts have been postulated for SLE, where antinuclear autoantibodies, including those directed against double-stranded DNA, have been detected up to 9.4 years before onset.^49^ Although we and others^50^ noted increasing autoantibody titres closer to disease onset, a titre threshold alone does not explain the transition from subclinical to clinical autoimmunity. In fact, some patients in our cohort had AQP4-IgG titres >1:1000 but did not develop clinically manifest NMOSD for more than 10 years. It is therefore possible that additional factors shape autoantibody pathogenicity, including epitope specificity, antibody affinity, and post-translational modifications such as glycosylation, that may evolve over time.^51,52^

In addition to autoantibody effector functions, clinical onset of NMOSD is thought to depend on the entry of AQP4-IgG into the CNS. NET formation may contribute to BBB disruption in NMOSD.^53^ Similarly, anti-TRIM21 autoantibodies from NMOSD patients have been shown to induce BBB dysfunction *in vitro.*^54^ Also known as Ro52/SSA autoantibodies, they are associated with SjD and SLE as well as higher disease activity in NMOSD.^55^ Interestingly, we observed lower levels of circulating TRIM21 in NMOSD cases compared to HC, suggesting antigen capture by anti-TRIM21 autoantibodies, as previously implicated.^41^ Interaction between TRIM21 and anti-TRIM21 autoantibodies was recently shown to perpetuate inflammation and promote (auto)antigen presentation through the formation of large immune complexes,^56^ warranting further investigation into how TRIM21 may contribute to NMOSD, SjD, and SLE pathogenesis.

Finally, we identified a 39-protein panel that differentiated preclinical NMOSD from HC in our sample set and warrants validation in independent cohorts. Although population-based screening is not feasible for a rare disease like NMOSD, this classifier may help identify individuals at risk among patients with related autoimmune conditions, enabling targeted AQP4-IgG screening. Our study further shows that preclinical NMOSD is distinct from the MS prodrome, where neuronal and myelin injury can be detected 6-7 years before onset,^11^ suggesting a potential window for preventive interventions before irreversible CNS damage occurs.

This study has limitations that are inherent to the DoDSR. First, the U.S. military population is predominantly male, whereas NMOSD disproportionately affects women (∼85:15 female-to-male ratio).^23^ Although our cohort still exhibited a female predominance with a 70:30 female-to-male ratio, a sex-related bias cannot be excluded. Second, to protect participant privacy, age was available only in categorical form, limiting the ability to adjust for age-related effects on proteins such as NEFL. Similarly, information on the exact timing of immunosuppressive therapy relative to post-onset sampling was unavailable. Another limitation is the variability in sampling intervals. Consecutive samples were often separated by several years, such that biological events (e.g. precise time of seroconversion) likely occurred earlier than observed. Regarding potential technical confounders, prolonged sample storage may have compromised protein stability. To account for this, normalized protein expression (NPX) composite scores, reflecting overall signal intensity, were incorporated into the statistical models, as described previously.^11^ For the machine learning analysis, limited sample size precluded independent model training and testing. Accordingly, performance estimates should be considered exploratory, pending validation in independent cohorts.

Collectively, this study provides insight into the preclinical evolution of NMOSD over a period of ∼24 years before onset. We demonstrate that innate immune dysregulation precedes AQP4-IgG seroconversion and CNS injury by several years and define a multi-hit model that extends current disease concepts. Independent *in vitro* and *in silico* analyses supported a potential role for CMV in NMOSD development. These findings provide a framework for understanding how immunological and environmental factors may converge during the earliest stages of autoimmunity, with implications for antibody-mediated diseases beyond NMOSD.

## Methods

### DoDSR cohort

NMOSD cases were identified among active-duty military personnel using the Defense Medical Surveillance System by International Classification of Diseases codes (ICD-9 241.0 and ICD-10 G36.0). Clinical records were reviewed and diagnoses were confirmed using the 2015 International Panel for NMO Diagnosis diagnostic criteria.^57^ Cases with ≥1 DoDSR specimen available were identified and up to four serum samples were obtained from each case as available: closest to disease onset, earliest sample available, and up to 2 samples in between these times. Each case was matched 1:1 by age, sex, and dates of collection of serum samples with HC. Chart reviews conducted by neurologists captured age at onset (recorded as a categorical variable due to data privacy restrictions), sex, presenting syndrome, co-existing autoimmunity, and immunotherapy exposure (**Table 1** and **Supplemental Table 4**). Based on retrospective CBA results, patients were classified as confirmed AQP4-IgG+ NMOSD, suspected seronegative NMOSD, MOGAD, or having an unclear autoantibody status. Only patients with confirmed AQP4-IgG+ or suspected seronegative NMOSD were included in the analysis (**Supplemental Figure 13**).

### Autoantibody testing

AQP4-IgG and MOG-IgG testing was performed on each NMOSD case and HC using a fixed CBA (Euroimmun, Lübeck, Germany) at ARUP Laboratories (Salt Lake City, UT). For NMOSD cases, testing began with the specimen closest to clinical onset or matched timepoint. If positive, progressively earlier samples were tested until a seronegative result was obtained or no prior specimen was available. For HC, AQP4-IgG and MOG-IgG testing was performed on the most recent serum sample, all samples tested negative. For all samples, autoantibody screening at an initial dilution of 1:10 was performed; if positive, additional dilutions were conducted at 1:4. The technician reading the CBA was blinded to the study group.

### Time interval definitions

For each NMOSD case included in our analysis, a median of three samples (range 2-4) was available. For each HC, a reference onset time was assigned based on the matched NMOSD patient. To assess protein changes across different time intervals during the preclinical phase and after disease onset, we evaluated multiple time-binning strategies to maximize the number of patients contributing one sample per interval. Based on this approach, the following temporal intervals were defined: samples collected within 90 days prior to onset or after were assigned to the “onset” timepoint; samples collected between 90 and 1,624 days before onset were categorized as “≤ 4.5 years before onset” (T1); samples between 1,625 and 2,749 days before onset as “4.5 - 7.5 years before onset” (T2); and samples collected ≥ 2,750 days before onset as “> 7.5 years before onset” or the “earliest timepoint” (T3). For duplicate samples per time intervals and patient (n = 17 HC and n = 9 NMOSD cases), samples were merged into a single mean NPX values. A summary of the samples per timepoint is outlined in **Supplemental Figure 2.**

### Proteomics proximity extension assays

Serum samples were centrifuged at 10,000 × g for 5 minutes. Subsequently, 50 µL aliquots were transferred into 96-well plates and analyzed, together with sample, plate, and negative controls, per the manufacturer’s protocol. Protein measurements were performed using the Olink Explore HT proximity extension assay, covering 5,401 proteins. Sequencing was carried out on an Illumina NovaSeqX platform. Each assay was normalized against five inter-plate controls. Intra- and inter-run coefficients of variation (CVs) were determined using three control samples included on each plate. The mean intra-run and inter-run CVs were 16.68% [10.25 - 28.61] and 22.53% [13.27 - 38.89], respectively. Sample quality was ensured using three internal spike-in controls to monitor the incubation, extension, and amplification steps. Preprocessing of proteomic data and quality control (QC) was conducted using the Olinkanalyze (v.4.5.0) package.^58^ Sample outliers were detected using principal component analysis (PCA), defined as samples exceeding ±3 standard deviations (SD) from the mean aggregated NPX values. Using this approach, only 9 of 313 samples (2.88%) were classified as outliers and excluded from the analysis (2 from NMOSD patients and 7 from HC). Samples/assays with QC warnings were excluded from the analysis. For all assays, relative protein abundance was reported as NPX values. As an additional QC step, we assessed technical agreement between Olink and Simoa platforms for NEFL and GFAP. Simoa assays (Quanterix Corp, Billerica, MA) were performed at Frontage Laboratories (Exton, PA). Specifically, the Simoa GFAP Discovery kit and NF-Light assay were employed in a 2-step immunoassay to quantify analyte concentration according to manufacturer specifications. We then fitted linear mixed-effects models (R package *lmerTest,* v.3.2.0)^59^ with Olink NPX as the dependent variable, log2-transformed Simoa concentration as a fixed effect, and a patient-specific random intercept (NPX ∼ log2(Simoa) + (1 | patientID)). We report the fixed-effect slope (β) with its standard error and p-value, alongside marginal (R²m) and conditional (R²c) coefficients, indicating a significant alignment between results generated using both methods (**Supplemental Figure 16**).

### Proteomics data analysis and statistics

Statistical analyses for proteomics were performed in R (v.4.5.2; 2025-10-31) and R Studio (v.2026.04.0+526). Group comparisons were performed using linear models implemented in the *limma* package (v.3.66.0).^60^ For each assay, individual NPX z-scores exceeding ± 3 from the respective mean were excluded from the model to reduce the effect of outliers. To account for any potential technical / pre-analytical variability affecting individual samples, as well as variance related to storage time, a composite NPX score (Sample_NPX_) was computed as the mean NPX across all protein assays per sample. In all group comparisons, Sample_NPX_, categorical age at collection (≥ 40 years; yes/no), and biological sex were included as covariates. For longitudinal comparisons with multiple samples per patient, we corrected for repeated measures using *duplicateCorrelation()* in *limma*. For *limma* comparisons including one sample per patient, lmFit with robust estimation was used. Log_2_FC and two-sided p-values for DEPs were reported, with an FDR of 5% (Benjamini-Hochberg procedure).^61^

In addition to the unbiased approach, we investigated a selection of proteins previously associated with NMOSD in a hypothesis-driven analysis: NEFL, GFAP, CXCL13, BCMA, BAFF/BAFF-R, the complement components C1r/s, C3, C4b, C5, C7, C8b, C9 and markers of Th17 differentiation (IL6, IL17A, IL23R).^17,29,40,51^ NPX values were visualized using boxplots and nominal p-values from *limma* analysis were reported. Linear mixed-effect models (R package *lmerTest,* v.3.2.0)^59^ were used to assess longitudinal changes in AQP4-IgG titres of patients with NMOSD.

Proteomic pathway analysis was conducted using preranked Gene Set Enrichment Analysis (GSEA) implemented in the R package *fgsea* (v1.36.2). All quantified proteins were ranked by moderated t-statistics derived from *limma* comparisons. Enrichment was assessed against Reactome pathways and Gene Ontology Biological Process (GO:BP) gene sets for *Homo sapiens*, retrieved via *msigdbr* (v25.1.1).^18,19^ Gene sets with 10-500 members were retained, and pathways were restricted to their intersection with the 5,401 proteins measured on the Olink panel. P-values from *fgsea* were FDR-adjusted,^61^ with results reported at FDR of 10% alongside Normalized Enrichment Scores (NES).

### Machine learning classifier

Onset samples from AQP4-IgG+ NMOSD patients and HC were included to build a supervised machine learning classifier. First, k-nearest neighbor imputation (k = 10; package *impute*, v.1.84.0) was used to impute missing values. We then applied Boruta feature selection (package *Boruta*, v.9.0.0) to identify the most important features (= proteins) for distinguishing NMOSD from HC. To assess selection stability, Boruta was repeated 20 times with different random seeds.^62^ Proteins confirmed in ≥ 60% of runs were retained for downstream model training with k-fold cross validation. To that end, model configurations were first tuned via a grid search over different alpha levels (α = 0, 0.1, 0.3, 0.5, 0.7, 0.9, 1.0), spanning Ridge (α = 0), Elastic Net (0 < α < 1), and LASSO (α = 1). For each α, the regularization strength λ was selected internally by *glmnet* (v4.1-10) using both lambda.min and lambda.1se criteria. Each (α, λ) configuration was then evaluated across 100 repeats of 5-fold cross-validation, with mean AUC as the selection metric. Ridge regression (α = 0, lambda.min) performed on the protein panel identified by Boruta feature selection achieved the highest mean AUC and was retained as the final model. Statistical significance was assessed by a label permutation test on selected features (1,000 permutations; 50 repeats of 5-fold CV per permutation), with the p value defined as the proportion of null AUCs ≥ the observed mean AUC. Given the established role of GFAP in symptomatic NMOSD, we also performed a sensitivity analysis comparing model performance with and without GFAP added to the Boruta-selected panel. ROC curves and AUCs were derived by averaging fold-wise ROC curves across the 5 CV folds within each repeat, then summarizing across 100 repeats. The final ROC curve was constructed from the pointwise mean sensitivity on a common specificity grid. We report the mean AUC across 100 repeats ± SD. Finally, the mean coefficient vector was computed as the row-wise average across 100 repeats. As an independent validation cohort was unavailable, the mean absolute coefficients and intercept were then applied to preclinical samples (T1-T3) of NMOSD cases and HC, with AUC and 95% confidence intervals (DeLong method)^63^ calculated per timepoint. The same approach was applied to distinguish post-onset AQP4-IgG+ from suspected seronegative NMOSD cases.

### GWAS

We performed GWAS in the UKB and AoU cohorts. In UKB, controls were first matched 100:1 to the broad ICD-10 G36 code by age, sex, and assessment center by nearest-neighbor matching, restricted to self-reported White participants. Cases were subsequently narrowed to an explicit G36.0 code (neuromyelitis optica). In AoU, cases were identified from electronic health records using ICD-10 G36.0, and controls matched 1:100 to cases by age, sex, and genetic ancestry. In both cohorts, cryptic relatedness was assessed with the KING-robust estimator (PLINK2), and one individual from each pair with kinship > 0.0442 (third-degree) was iteratively removed. Population structure was captured by the top 20 (UKB) or top 10 (AoU) principal components computed on LD-pruned autosomal variants. Association testing was restricted to common variants (MAF > 1%) using REGENIE v3.2.1 with Firth correction adjusting for age, sex, and the corresponding PCs. We combined and harmonized the UKB and AoU results with three publicly available GWAS in European^21^, Asian^22^, and pan-ancestry^23^ cohorts (**Supplemental Table 2**). All summary statistics were harmonized to GRCh37/hg19 and palindromic variants (A/T, C/G) were excluded. Random-effects meta-analysis was performed in GWAMA v2.2.2^64^. Gene-level associations for CXCL1 were tested with MAGMA^20^ (variants within 50 kb; combined 1000 Genomes Phase 3 LD reference), applied to the meta-analyses as well as to each cohort individually.

### VirScan

To profile antiviral antibody repertoires, we performed VirScan,^25^ a high-throughput PhIP-Seq technology with an expanded viral peptide library of more than 480,000 synthetic peptides.^65^ Samples were run in duplicates and processed as described previously.^66^ Briefly, deep-well 2 mL round-bottom 96-well plates were blocked overnight at 4°C using 3% bovine serum albumin (BSA) in Tris-Buffered Saline with Tween-20 to reduce non-specific binding. The blocking buffer was removed and replaced with 500 µL of the phage library (10^10^ plaque-forming units per milliliter), alongside 1 µL of human serum diluted 1:1 in storage buffer (phosphate-buffered saline supplemented with 0.04% sodium azide, 40% glycerol, and 40 mM HEPES) for immunoprecipitation. Plates were incubated overnight at 4°C on a rocking platform to facilitate antibody-phage interactions. Protein A and G magnetic beads (Thermo Fisher Scientific, 10002D and 10004D) were washed in TNP-40 buffer and added to each well to capture immune complexes. After 1 hour at 4°C, beads were transferred to filter plates compatible with vacuum manifolds and subjected to 5 washes with RIPA buffer. Between each wash, beads were incubated for 5 minutes at 4°C with rocking. 150 µL LB-Carb was added to the immunoprecipitated phage and amplified in 550 µL log-phase *Escherichia coli* strain BLT5403 (OD₆₀₀ = 0.3-0.6). After 2 hours of shaking at 1200 rpm, NaCl was added to a final concentration of 0.5 M to facilitate full lysis. Lysates were centrifuged at 3500 rpm for 1 hour. The supernatant was transferred to a separate BSA-blocked round-bottom plate, and a second round of immunoprecipitation and amplification was performed to enrich specific antibody-bound phage. Final lysates were centrifuged and stored at 4°C for subsequent library preparation. Phage DNA was barcoded and amplified using Phusion PCR (18 cycles), then subjected to sequencing on an Illumina NovaSeq platform, yielding approximately 1 million reads per sample.

VirScan data analysis was performed in R (v.4.5.2; 2025-10-31) and R Studio (v.2026.1.1.403). Peptides were annotated by viral family, genus, and species. Downstream analyses were restricted to human-infecting viruses. For each replicate, reads per 100,000 (RPK) was assessed. Bead RPK values were used to calculate z-scores and log_2_FC relative to beads. Before group comparisons, a group-agnostic threshold-above-beads filter was applied, where a peptide was considered positive only if both replicates satisfied three bead-based criteria: minimum raw RPK > 10, fold change > 10, and z-score > 3 in at least two unique patients. For peptides meeting these thresholds, RPKs and log_2_FC values were averaged across replicates. Peptide-level group comparisons were performed on the filtered peptide universe using *limma* package (v3.66.0),^60^ applied to the sample-level mean log_2_FC versus beads. All tests were two-sided; within-patient correlation was estimated using *duplicateCorrelation* for paired measurements. For paired pre/post-seroconversion analysis, threshold positivity was collapsed to paired patient-level status and tested using an exact binomial test based on discordant pairs. For unpaired comparisons, threshold positivity was collapsed to the patient level and compared between groups using Fisher exact tests. For significant peptides at FDR 5%,^61^ we further assessed whether reactivity signals were enriched compared to a HC background using HC samples from similar timepoints.

### SPOKE analysis

Network construction and statistical analyses were performed in Python 3.12 (igraph 0.10, numpy, pandas, scipy.stats, statsmodels.stats.multitest); figure assembly in R 4.4 (ggplot2, ggraph, patchwork, ggsignif). First, we identified DEPs at FDR 5% from three *limma* group comparisons (pre-seroconversion/pre-onset, post-seroconversion/pre-onset, and post-onset NMOSD vs. HC). We then tested whether the NMOSD plasma proteome shows preferential proximity to bacterial or viral organisms in the human interactome. We used a SPOKE subgraph of 223,202 nodes and 4,082,356 edges, containing eleven node types: human proteins, genes, organisms, diseases, GO biological processes, GO molecular functions, GO cellular component, Reactome pathway, anatomy, cell type, non-human proteins.^28^ DEP UniProt accessions were used as the seed set *S* of a Personalised PageRank random walk ^67^ on the graph (igraph implementation ^68^; damping factor *α* = 0.85, restart probability 1 − *α* = 0.15). The stationary PPR distribution ***π*** satisfies

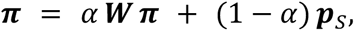

where ***W*** is the column-stochastic weighted transition matrix and ***p****_S_* is the seed-restart distribution (uniform on *S*, zero elsewhere). To control for the dependence of PPR mass on node degree, a degree-matched permutation null was generated by drawing 500 random seed sets in which each true seed was replaced by a protein sampled from a degree-matched pool (±20% of the seed’s degree, minimum pool size 10). For each organism node *i*, the empirical z-score and one-sided p-value were computed as

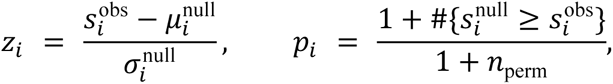

and p-values were Benjamini-Hochberg-adjusted within node type.^61^ Organism-level significance is reported at FDR 5%. Each organism node was assigned to a superkingdom by hierarchical lookup against the NCBI species taxonomy,^69^ falling back to a curated alias map that reconciles legacy NCBI and modern ICTV nomenclatures (e.g. *Human herpesvirus 5* ↔ *Cytomegalovirus humanbeta5*; *Borrelia* ↔ *Borreliella*). This procedure assigned a kingdom to 424 of the 597 organism nodes, i.e. 166 Viruses and 258 Bacteria. The kingdom-level effect size was quantified by Cliff’s δ,^70^

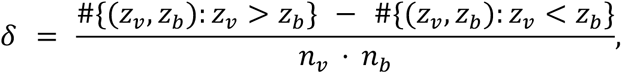

where *z_v_*, *z_b_* are PPR z-scores of viral and bacterial organisms, respectively (range −1 to +1; negative values indicate bacteria-biased proximity). A 95% confidence interval was obtained from 5,000 bootstrap resamples (sampling with replacement within each kingdom). The null hypothesis of equal kingdom distributions was tested by 5,000 label-permutation iterations on Cliff’s δ (two-sided empirical p-value); the mean-difference *μ_V_* − *μ_B_* was tested in parallel under the same scheme.

For visualization, strain-level entries within each comparison were collapsed to the species level by retaining the maximum |*z*| per (species, kingdom) pair after stripping strain, subspecies, and alphanumeric suffix tokens. To resolve the molecular content of the top organism hits, a directed sub-graph was extracted from SPOKE following the metapath *Organism → pathogen protein → human target → DEP* by traversing three edge layers: organism → pathogen-protein edges (5,237), pathogen–host PPI edges (92,656) and human–human PPI edges (1,498,917). Direct overlap (DEP equal to a human target) and one-hop extension (DEP equal to a 1-hop human neighbor of a target) were both retained; each sub-graph was pruned to the most strongly regulated DEPs (by |log_2_FC|) and the most-connected human hubs, then laid out by stress minimization with ggraph.

## Data availability

The proteomic dataset and related clinical data will be made available via Dryad upon journal publication.

## Code availability

No original tool was developed for this paper. All the applied statistical methods are publicly available and referenced in the paper.

## Ethical approval

This study was approved by the University of Utah Institutional Review Board (IRB#00090978) and WRNMMC-2017-0046 EIRB. Participant consent was waived.

## Acknowledgements and funding statement

This study was supported by funding from Alexion/AstraZeneca Rare Disease (SRA Proposal 200157), the Western Institute for Veterans Research (both to TLS), and the Sumaira Foundation (to SLC). AA was supported by Career Transition Award from the National Multiple Sclerosis Society (NMSS; TA-2404-43033) and K23 award from the National Institute of Neurological Disorders and Stroke (K23NS140543). MRW was supported by the Westridge Foundation, and Tim and Laura O’Shaughnessy. LMJ was supported by the Deutsche Forschungsgemeinschaft (DFG, MU 5558/1-1) and the Else Kröner-Fresenius Stiftung (2023_EKEA.197). LMJ is furthermore a participant in the BIH Charité Clinician Scientist Program funded by the Charité - Universitätsmedizin Berlin and the Berlin Institute of Health at Charité, Germany. DC is supported by the NMSS postdoctoral funding. GB is supported by a research fellowship FISM - Fondazione Italiana Sclerosi Multipla – cod. 2024/BR/005 and financed or co-financed with the ‘5 per mille’ public funding. We thank all patients and military personnel for their valuable contribution to this study.

## Author contribution statement

**LMJ:** Conception of the study, statistical analyses, data interpretation, drafting of the manuscript. **DC:** VirScan analyses, statistical analysis, drafting and revision of the manuscript. **GB:** SPOKE analyses, data analysis, drafting and revision of the manuscript. **YD:** GWAS analysis, drafting and revision of the manuscript. **MH:** Sample processing, proteomics assays, drafting and revision of the manuscript. **AMG:** GWAS analysis, drafting and revision of the manuscript. **CEM**: Data interpretation, drafting and revision of the manuscript. **RD**: VirScan analyses, statistical analysis, drafting and revision of the manuscript. **JWR:** Data acquisition, drafting and revision of the manuscript. **LKP:** autoantibody cell-based assays, drafting and revision of the manuscript. **KHW**: Conception of the study, drafting and revision of the manuscript. **BLD**: NEFL and GFAP assays, drafting and revision of the manuscript. **NC:** Data acquisition, drafting and revision of the manuscript. **DEH:** Data acquisition, drafting and revision of the manuscript. **KG:** Data acquisition, drafting and revision of the manuscript. **NT:** Data acquisition, r drafting and revision of the manuscript. **SRO:** Data acquisition, drafting and revision of the manuscript. **SLM:** Data and sample acquisition, drafting and revision of the manuscript. **RBC:** Data acquisition, drafting and revision of the manuscript. **VW:** Data acquisition, drafting and revision of the manuscript. **MMPS:** Data acquisition, drafting and revision of the manuscript. **SJP:** Conception of the study, drafting and revision of the manuscript. **BACC:** Data interpretation, drafting and revision of the manuscript. **SSZ**: Data interpretation, drafting and revision of the manuscript. **AJG:** Data interpretation, drafting and revision of the manuscript. **AH:** GWAS analysis, drafting and revision of the manuscript. **SEB:** SPOKE data interpretation, drafting and revision of the manuscript. **MRW:** Conception of the study, VirScan analyses, data interpretation, supervision, drafting and revision of the manuscript. **SLC:** Conception of the study, data acquisition, data interpretation, supervision, drafting and revision of the manuscript. **AA:** Conception of the study, funding, proteomics assays, statistical analysis, data interpretation, supervision, drafting and revision of the manuscript. **TLS:** Conception of the study, funding, data interpretation, supervision, drafting and revision of the manuscript. All authors reviewed and approved the final manuscript.

## Competing interest declaration

AMG is supported by the Stage Quebec program and the Canadian Institute for health Research (CIHR) doctoral scholarship, funding reference number: 210230. CEM served on an advisory board for Sanofi, the editorial review board for Evidence to Action, and is listed on a patent application related to autoimmune aquaporinopathy. CEM has been supported by a National Institutes of Health Mentored Clinician Scientist Research Career Development Award (K08AI199129), a Clinician Scientist Development Award (FAN-2107-38301) and a Career Transition Award (TA-2503-44476) both from the National Multiple Sclerosis Society, a Neuroscience Research Training Scholarship from the American Academy of Neurology, a Clinician-Scientist Award from the Weill Neurohub, the J.M. Hampton, Jr. Medical Research Fund from the UCSF Research Evaluation and Allocation Committee (REAC), and the C.D. Spangler Foundation. MMP serves on the editorial board for Journal of Neuroimaging; has received research funding from Biogen, Novartis, and Clene Nanomedicine; and has served on an advisory board for TG therapeutics. LKP has served on the advisory boards of Werfen and AliveDx, is head of clinical operations at ARUP laboratories, is associate editor at Clinical Biochemistry and LabQ, and is president of the Association of Medical Laboratory Immunologists. SJP has received personal compensation for consulting or serving on scientific advisory or data safety monitoring boards for Arialys, Astellas, F. Hoffmann-La Roche AG, Genentech, Sage Therapeutics, and UCB; his institution has received compensation for his serving as a consultant for Alexion, Amgen, Arialys, and Astellas; research support from Adimmune, Alexion, Amgen, F. Hoffmann-La Roche AG, and Genentech; and has 2 patents issued (8889102; application 12-678350; Neuromyelitis Optica Autoantibodies as a Marker for Neoplasia; and 9891219B2; application 12-573942; Methods for Treating Neuromyelitis Optica [NMO] by Administration of Eculizumab to an Individual that is Aquaporin-4 [AQP4]-IgG Autoantibody Positive), for which he has received royalties. AH has received speaker and/or consultancy fees from Alexion, Amgen, Biogen, Novartis, Pfizer, and Roche; compensation for serving on a scientific advisory board for Amgen, Novartis, and Roche; and research funding from Amgen and Roche, all unrelated to the present work, AH is further supported by a Clinical Research Scholarship (349722) and a Clinician-Researcher Establishment Grant (358114) from the Fonds de Recherche du Québec Santé (FRQS), the Bougie Family Young Investigator Award, as well as a Future Leaders in Canadian Brain Research Grant from the Brain Canada Foundation, AH is further funded by the Canada Brain Research Fund (CBRF) and by the Azrieli Foundation. AJG reports research support from F.Hoffmann-La Roche, NIH, DoD and NMSS as well as personal fees from Neurona, Borealis Bioscience and Pipeline Therapeutics and has served as an editor at JAMA Neurology. MRW receives unrelated research grant funding from Roche/Genentech, Novartis, and Kyverna Therapeutics and is a founder and board member of Delve Bio, Inc. He has done consulting for Pfizer, Vertex Pharmaceuticals, Ouro Medicines, Red Tree Ventures and Indapta Therapeutics; SLC reports serving as a local site investigator for the UCB-sponsored cosMOG trial and the SPHERES registry, SLC has received research funding paid to their institution from the NIH/NINDS, the Siegel Rare Neuroimmune Association, Amgen, Alexion/AstraZeneca Rare Disease, and Kyverna, SLC has served as a consultant or advisory board member for Alexion/AstraZeneca Rare Disease and Amgen, served on the data and safety monitoring board for Kyverna, and serves on the Medical Advisory Boards of the Sumaira Foundation for NMO and the Autoimmune Encephalitis Alliance, SLC is also Editor of the *Neurology* Podcast and *Neurology Minute* and serves on the Editorial Board of *Neurology: Neuroimmunology & Neuroinflammation*; AA received research grants from the National Institute of Health, the National Multiple Sclerosis Society, the German Multiple Sclerosis Society, the Department of Defense, UCSF Weill Institute for Neurosciences, and UCSF Sandler Program for Breakthrough Biomedical Research, he also received speaker/ consultation fees from Genentech/Roche and Sanofi; TLS has received research funding from Alexion/AstraZeneca and the Western Institute for Veterans Research to conduct this study, she has received compensation for serving on scientific advisory boards for Amgen, EMD Serono, and UCB. All other authors report no conflicts related to this study.

