## Supplemental_Material for "Innate Immune Dysregulation Precedes Clinical Onset in Neuromyelitis Optica"

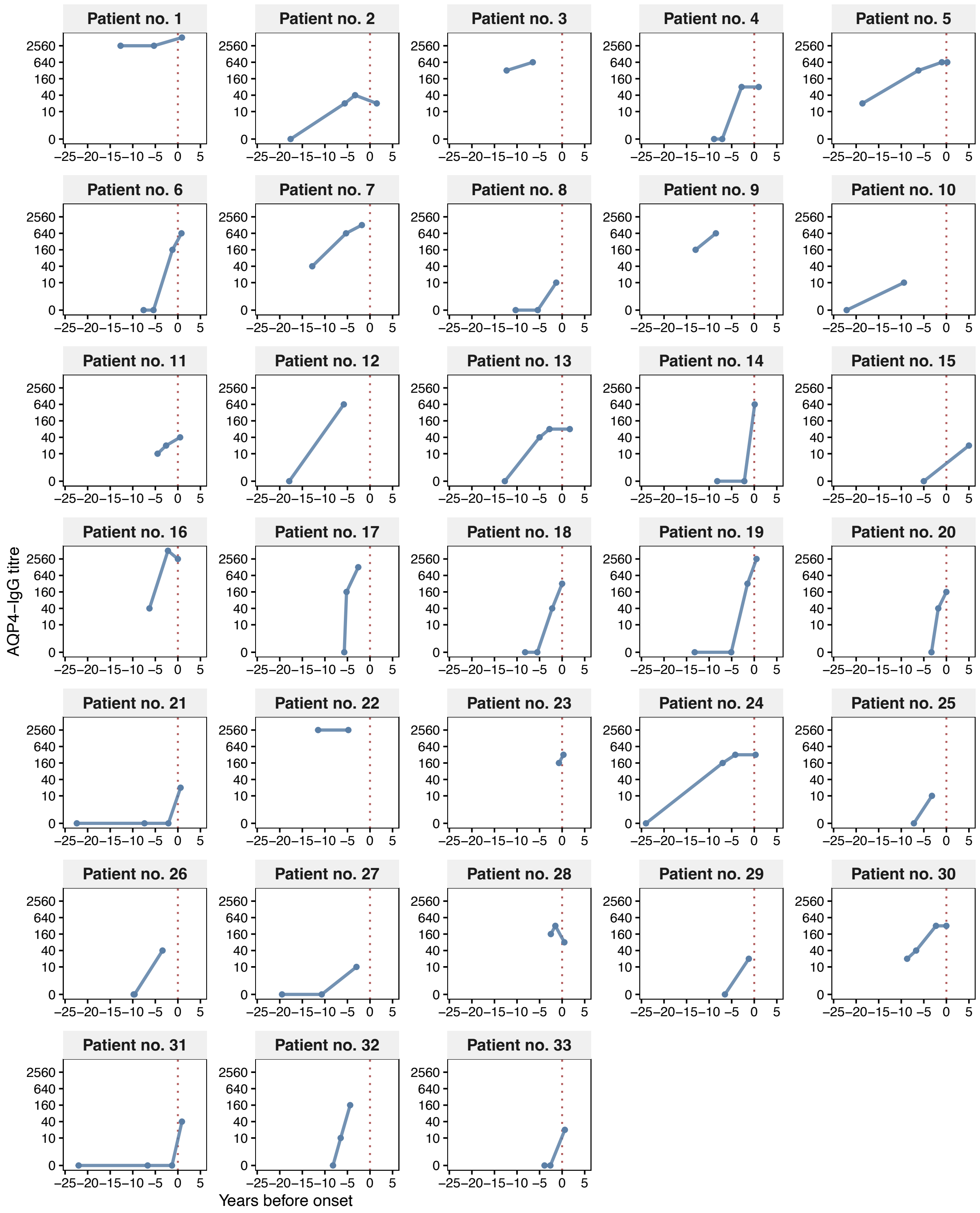


**Supplemental Figure 1. AQP4-IgG titre trajectories in patients with NMOSD.** Y axis shows $\log_2$ AQP4-IgG titre alongside zero values, x axis shows years before clinical onset. The red dotted line depicts clinical onset of NMOSD.

**Supplemental Table 1. List of differentially expressed serum proteins between patients with AQP4-IgG+ NMOSD and healthy controls including all longitudinal samples (FDR of 5%).**

| Assay | UniProt ID | $Log_2$ fold change | Nominal *P*-value | Adjusted *P*-value |
|---|---|---|---|---|
| **CXCL1** | P09341 | 1.16 | <0.001 | <0.001 |
| **CXCL3** | P19876 | 1.32 | <0.001 | <0.001 |
| **CXCL5** | P42830 | 0.97 | <0.001 | <0.001 |
| **CCL24** | O00175 | -0.74 | <0.001 | <0.001 |
| **CXCL11** | O14625 | 0.9 | <0.001 | <0.001 |
| **CLEC4D** | Q8WXI8 | -0.78 | <0.001 | <0.001 |
| **CPM** | P14384 | -0.37 | <0.001 | <0.001 |
| **ENPP1** | P22413 | -0.68 | <0.001 | <0.001 |
| **BPIFA2** | Q96DR5 | 1.06 | <0.001 | <0.001 |
| **KLRB1** | Q12918 | -0.34 | <0.001 | <0.001 |
| **CTSW** | P56202 | -0.66 | <0.001 | <0.001 |
| **CD207** | Q9UJ71 | -0.41 | <0.001 | <0.001 |
| **SRPX** | P78539 | 0.43 | <0.001 | <0.001 |
| **VNN 2.00** | O95498 | -0.39 | <0.001 | <0.001 |
| **RAET1L_ULBP2** | Q5VY80_Q9BZM5 | -0.34 | <0.001 | <0.001 |
| **KLK15** | Q9H2R5 | -0.39 | <0.001 | <0.001 |
| **PRG3** | Q9Y2Y8 | -0.5 | <0.001 | <0.001 |
| **CXCL6** | P80162 | 0.79 | <0.001 | <0.001 |
| **CCL15** | Q16663 | 0.48 | <0.001 | <0.001 |
| **CCL19** | Q99731 | 1 | <0.001 | <0.001 |
| **C7** | P10643 | 0.31 | <0.001 | <0.001 |
| **LRFN2** | Q9ULH4 | 0.73 | <0.001 | <0.001 |
| **ITGA11** | Q9UKX5 | -0.32 | <0.001 | <0.001 |
| **GPC5** | P78333 | -0.4 | <0.001 | <0.001 |
| **LPO** | P22079 | 0.58 | <0.001 | <0.001 |
| **TFRC** | P02786 | 0.29 | <0.001 | 0.001 |
| **SLAMF7** | Q9NQ25 | 0.52 | <0.001 | 0.001 |
| **OMD** | Q99983 | 0.46 | <0.001 | 0.002 |
| **CST5** | P28325 | 0.43 | <0.001 | 0.002 |
| **SFTPD** | P35247 | 0.5 | <0.001 | 0.002 |
| **FUCA1** | P04066 | 0.41 | <0.001 | 0.002 |
| **C4BPB** | P20851 | 0.33 | <0.001 | 0.002 |
| **GSTM4** | Q03013 | 1.09 | <0.001 | 0.002 |
| **AMY1_AMY2** | P04746_P0DTE7_P0DTE8_P0DUB6_P19961 | 0.34 | <0.001 | 0.002 |
| **FOLR3** | P41439 | 1.9 | <0.001 | 0.003 |
| **NOTCH2** | Q04721 | 0.34 | <0.001 | 0.003 |
| **CTSH** | P09668 | 0.57 | <0.001 | 0.003 |
| **CXCL13** | O43927 | 0.61 | <0.001 | 0.003 |
| **SSC4D** | Q8WTU2 | -0.83 | <0.001 | 0.004 |

| Assay | UniProt ID | Log$_2$ fold change | Nominal *P*-value | Adjusted *P*-value |
|---|---|---|---|---|
| **ADAM22** | Q9P0K1 | -0.35 | <0.001 | 0.004 |
| **SMPDL3A** | Q92484 | 0.45 | <0.001 | 0.005 |
| **MCEMP1** | Q8IX19 | -0.73 | <0.001 | 0.005 |
| **CLEC11A** | Q9Y240 | 0.3 | <0.001 | 0.005 |
| **PRR4** | Q16378 | 0.49 | <0.001 | 0.006 |
| **AHNAK** | Q09666 | -0.37 | <0.001 | 0.007 |
| **CLSTN3** | Q9BQT9 | -0.28 | <0.001 | 0.007 |
| **ICAM5** | Q9UMF0 | -0.31 | <0.001 | 0.007 |
| **CNTN3** | Q9P232 | -0.19 | <0.001 | 0.007 |
| **LBP** | P18428 | 0.52 | <0.001 | 0.007 |
| **DSG4** | Q86SJ6 | -0.33 | <0.001 | 0.008 |
| **LRIG1** | Q96JA1 | 0.29 | <0.001 | 0.008 |
| **FMNL1** | O95466 | -0.58 | <0.001 | 0.008 |
| **SPINK7** | P58062 | 0.57 | <0.001 | 0.009 |
| **GFRAL** | Q6UXV0 | -0.46 | <0.001 | 0.010 |
| **HPCAL4** | Q9UM19 | 0.4 | <0.001 | 0.010 |
| **CLMP** | Q9H6B4 | -0.23 | <0.001 | 0.011 |
| **CHI3L1** | P36222 | -0.33 | <0.001 | 0.011 |
| **HOMER3** | Q9NSC5 | -0.52 | <0.001 | 0.011 |
| **NCR1** | O76036 | -0.23 | <0.001 | 0.011 |
| **ICAM3** | P32942 | 0.21 | <0.001 | 0.011 |
| **RBP7** | Q96R05 | -0.57 | <0.001 | 0.012 |
| **APOF** | Q13790 | -0.35 | <0.001 | 0.012 |
| **RRM2** | P31350 | 0.41 | <0.001 | 0.012 |
| **IL15** | P40933 | 0.24 | <0.001 | 0.012 |
| **IL32** | P24001 | 0.34 | <0.001 | 0.012 |
| **ENDOU** | P21128 | -0.28 | <0.001 | 0.013 |
| **COL24A1** | Q17RW2 | -0.28 | <0.001 | 0.013 |
| **SCGB3A2** | Q96PL1 | 0.62 | <0.001 | 0.013 |
| **CXCL16** | Q9H2A7 | 0.22 | <0.001 | 0.013 |
| **SERPINB4** | P48594 | -0.39 | <0.001 | 0.013 |
| **HSD11B1** | P28845 | 0.28 | <0.001 | 0.013 |
| **GSR** | P00390 | 0.18 | <0.001 | 0.014 |
| **NCF2** | P19878 | -0.65 | <0.001 | 0.015 |
| **SEMA3G** | Q9NS98 | 0.24 | <0.001 | 0.015 |
| **GALNT5** | Q7Z7M9 | -0.35 | <0.001 | 0.015 |
| **CYB5R2** | Q6BCY4 | -0.56 | <0.001 | 0.015 |
| **IFNA1_IFNA13** | P01562 | 0.48 | <0.001 | 0.016 |
| **NRP2** | O60462 | 0.22 | <0.001 | 0.017 |
| **VIT** | Q6UXI7 | 0.3 | <0.001 | 0.017 |
| **CCL8** | P80075 | 0.39 | <0.001 | 0.017 |

| Assay | UniProt ID | Log$_2$ fold change | Nominal *P*-value | Adjusted *P*-value |
|---|---|---|---|---|
| **FCRL5** | Q96RD9 | 0.38 | <0.001 | 0.017 |
| **TRIM21** | P19474 | -0.52 | <0.001 | 0.017 |
| **FGFR4** | P22455 | -0.35 | <0.001 | 0.017 |
| **ANG** | P03950 | 0.21 | <0.001 | 0.018 |
| **MUC1** | P15941 | -0.38 | <0.001 | 0.018 |
| **FCAMR** | Q8WWV6 | 0.39 | <0.001 | 0.018 |
| **SLC39A5** | Q6ZMH5 | -0.31 | <0.001 | 0.020 |
| **COL2A1** | P02458 | -0.63 | <0.001 | 0.020 |
| **IL5RA** | Q01344 | 0.35 | <0.001 | 0.020 |
| **DBH** | P09172 | -0.43 | <0.001 | 0.021 |
| **STAB2** | Q8WWQ8 | -0.19 | <0.001 | 0.022 |
| **LGALS9** | O00182 | 0.23 | <0.001 | 0.023 |
| **NCF1C_NCF1** | A8MVU1_P14598 | -0.52 | <0.001 | 0.024 |
| **CD300LF** | Q8TDQ1 | -0.46 | <0.001 | 0.024 |
| **DAPK2** | Q9UIK4 | -0.54 | <0.001 | 0.024 |
| **CXCL10** | P02778 | 0.49 | <0.001 | 0.024 |
| **PRTG** | Q2VWP7 | -0.19 | <0.001 | 0.024 |
| **NME3** | Q13232 | 0.19 | <0.001 | 0.025 |
| **CD160** | O95971 | -0.26 | <0.001 | 0.025 |
| **SPINK4** | O60575 | 0.36 | <0.001 | 0.026 |
| **GPD1** | P21695 | -0.49 | <0.001 | 0.026 |
| **PTN** | P21246 | -0.9 | <0.001 | 0.027 |
| **CFHR4** | Q92496 | 0.18 | <0.001 | 0.027 |
| **GAS2L3** | Q86XJ1 | -0.66 | <0.001 | 0.031 |
| **CNDP1** | Q96KN2 | -0.33 | <0.001 | 0.031 |
| **TNFRSF11B** | O00300 | 0.24 | <0.001 | 0.032 |
| **ADH1B** | P00325 | -0.56 | <0.001 | 0.032 |
| **TNFSF15** | O95150 | 0.32 | <0.001 | 0.033 |
| **PODXL** | O00592 | -0.1 | <0.001 | 0.033 |
| **C1QTNF9** | P0C862 | -0.35 | <0.001 | 0.033 |
| **CCL5** | P13501 | 0.55 | <0.001 | 0.033 |
| **SELP** | P16109 | -0.3 | <0.001 | 0.033 |
| **NID2** | Q14112 | -0.4 | <0.001 | 0.034 |
| **SPINK5** | Q9NQ38 | -0.22 | <0.001 | 0.034 |
| **TFPI2** | P48307 | 0.33 | <0.001 | 0.036 |
| **NMNAT1** | Q9HAN9 | -0.56 | <0.001 | 0.036 |
| **VSTM4** | Q8IW00 | -0.54 | <0.001 | 0.037 |
| **PSTPIP1** | O43586 | -0.57 | <0.001 | 0.038 |
| **GHR** | P10912 | -0.25 | <0.001 | 0.038 |
| **SIGLEC1** | Q9BZZ2 | 0.3 | <0.001 | 0.038 |
| **ADGRE3** | Q9BY15 | -0.28 | <0.001 | 0.043 |

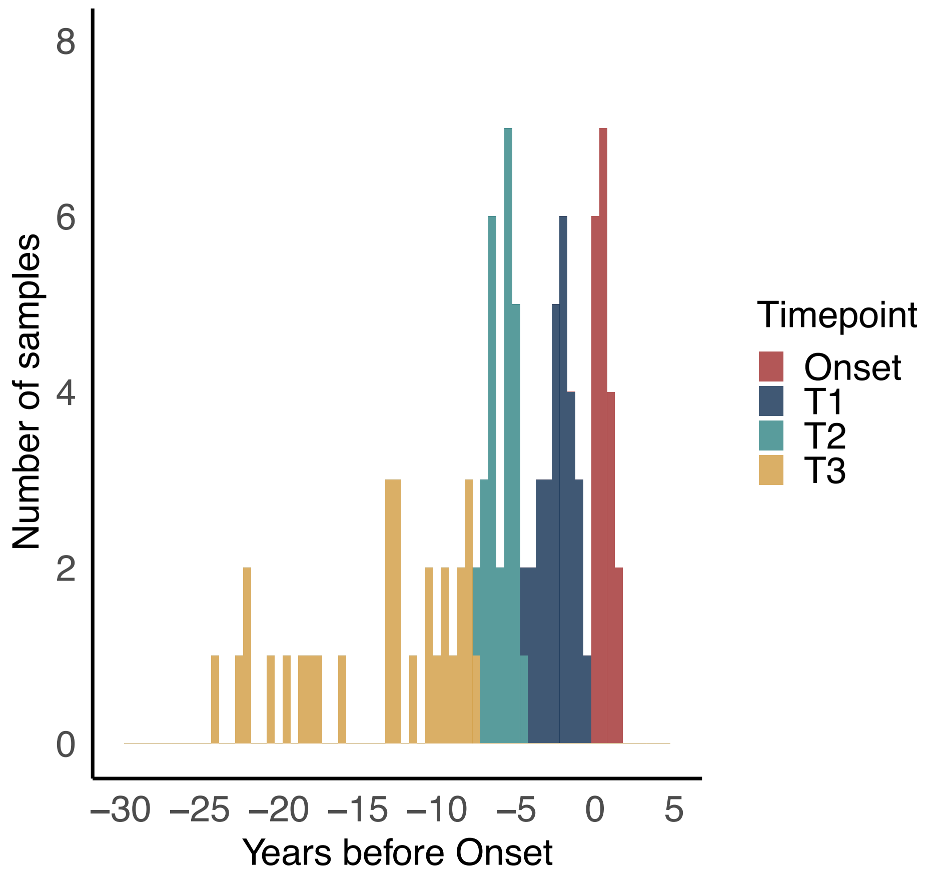


**Supplemental Figure 2. Sampling timepoints relative to clinical onset of NMOSD.** Histogram showing the distribution of samples from NMOSD patients relative to disease onset. The x-axis shows time (years before clinical onset), and y-axis shows the number of samples per 0.5-year interval. Colors indicate four time intervals: Onset (samples collected < 90 days before onset or after onset), T1 (samples collected between day -90 and day -1624; ~ < 4.5 years before onset), T2 (samples collected between day -1625 and -2749; ~ < 7.5 years before onset), and T3 (samples collected at day -2750 or earlier; ~ >7.5 years before onset).

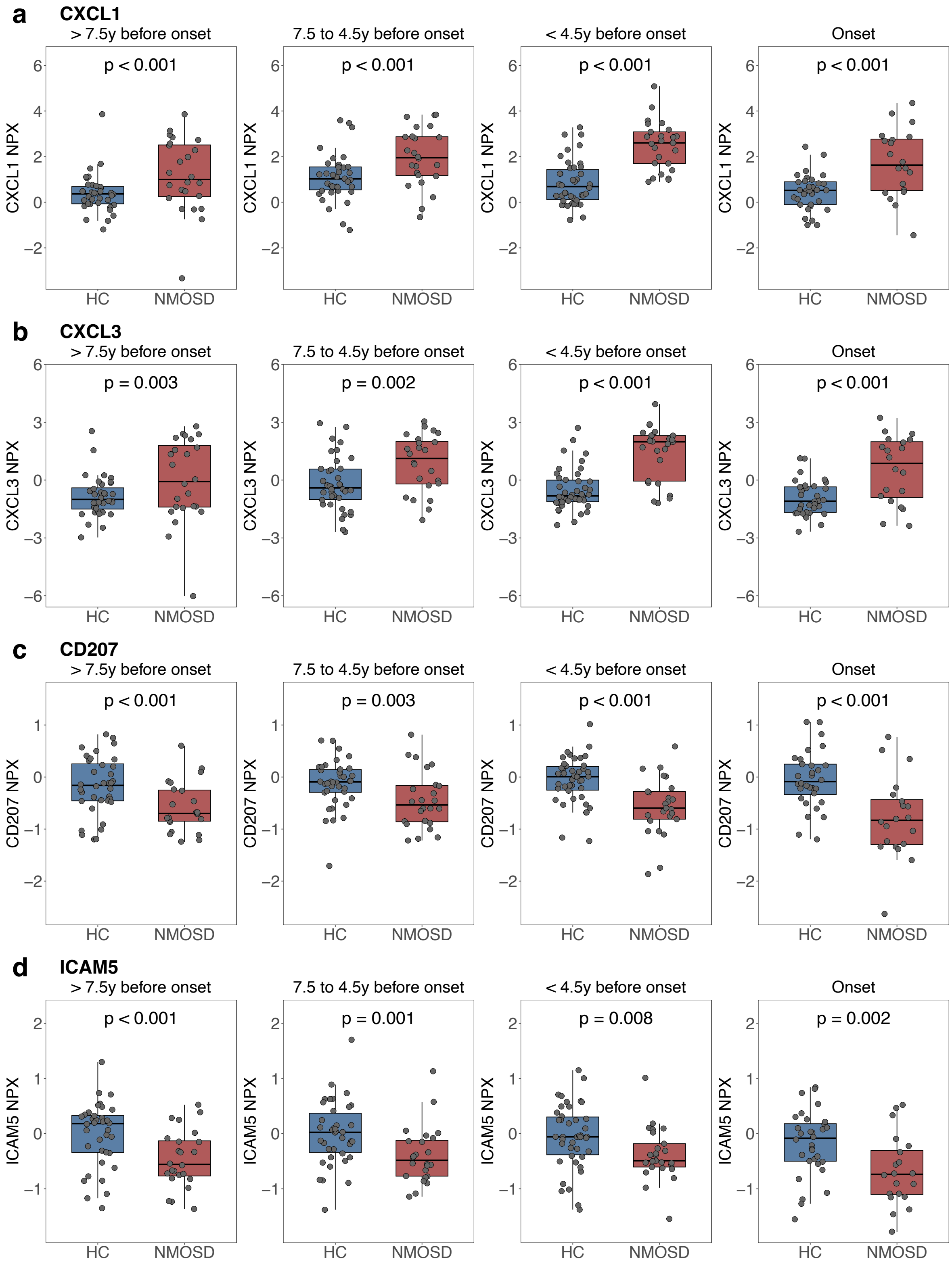


**Supplemental Figure 3. Abundance of consistently differentially expressed serum proteins between patients with NMOSD and HC across different time intervals.** P-value annotations display nominal p-values derived from *limma* comparisons, y; years. Boxplots display the median normalized protein expression (NPX) value (center line), interquartile range (box), and 1.5 × the interquartile range (whiskers). HC, healthy controls; NMOSD, neuromyelitis optica spectrum disorder.

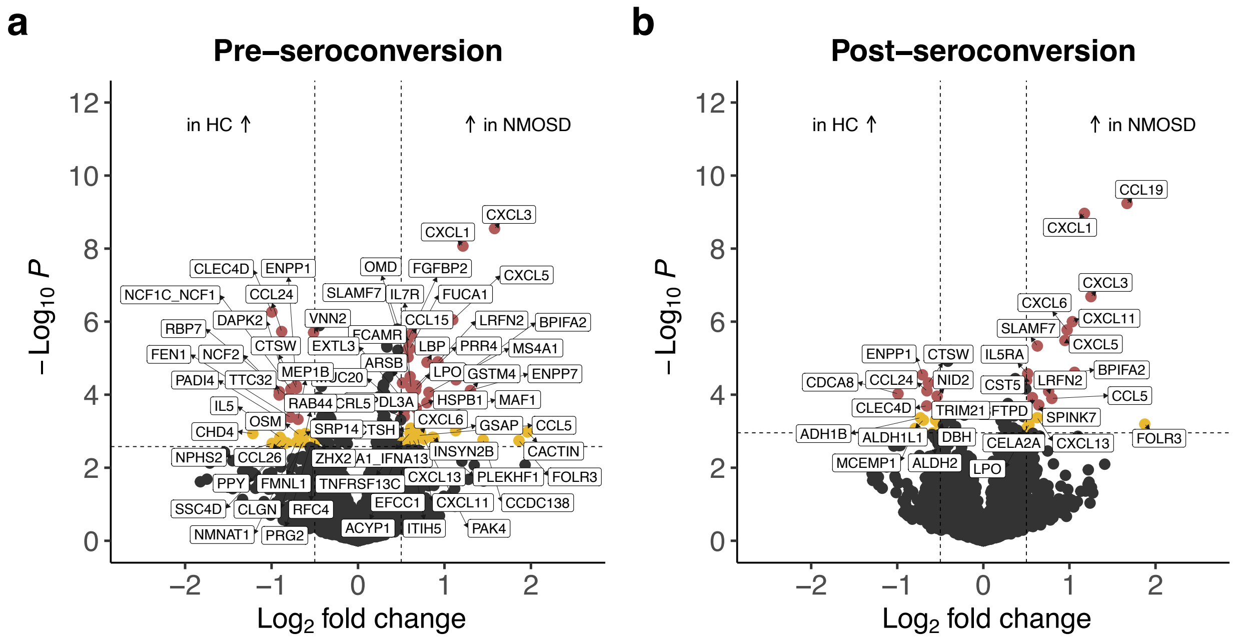


**Supplemental Figure 4. Differentially expressed serum proteins (DEPs) between patients with NMOSD and HC stratified by AQP4-IgG seroconversion status. (a)** Volcano plot showing DEPs between NMOSD cases, with samples collected before AQP4-IgG seroconversion (n = 35), and healthy controls (HC; n = 133). **(b)** Volcano plot showing DEPs between NMOSD cases, with samples collected after AQP4-IgG seroconversion but pre-onset (n = 45), and HC (n = 133). Proteins increased in HC are shown on the left, and proteins increased in NMOSD on the right. Significant DEPs at FDR 5% with a $\log_2$ fold change ($\log_2$FC) > 0.5 are highlighted in red and labeled. Significant DEPs at FDR 10% with a $\log_2$FC > 0.5 are shown in yellow and labeled. Vertical dashed lines indicate a $\log_2$FC threshold of > 0.5, horizontal dashed lines indicate an FDR threshold of 10%. HC, healthy controls; NMOSD, neuromyelitis optica spectrum disorder.

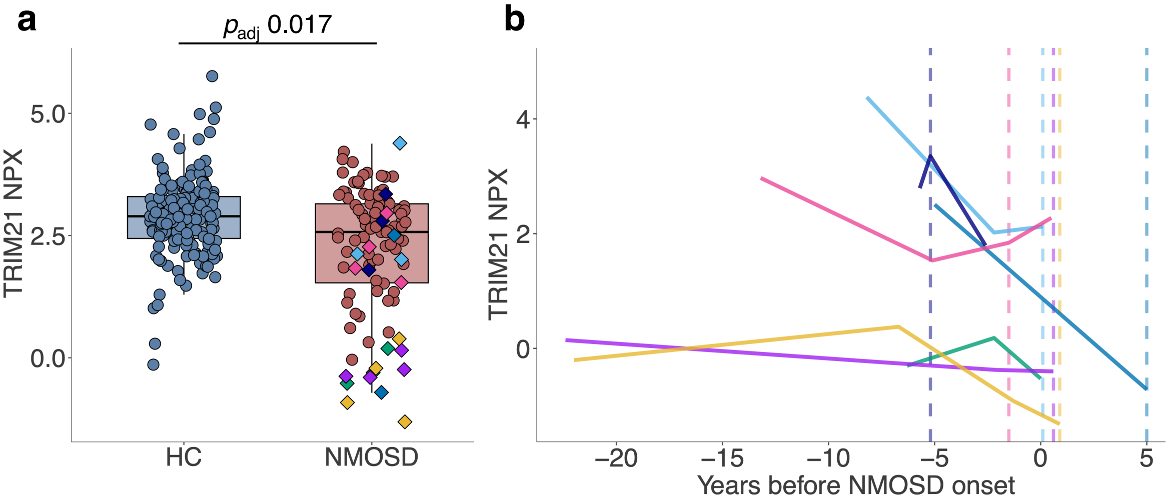


**Supplemental Figure 5. Abundance of TRIM21 in NMOSD. (a)** Comparison of TRIM21 normalized protein expression (NPX) values between NMOSD cases and healthy controls (HC), with individual data points representing all samples included in the study (pre- and post-onset). Boxplots display the median (center line), interquartile range (box), and 1.5 × the interquartile range (whiskers). P-values were calculated using a linear model with *limma* (FDR 5%). Colored diamonds indicate samples from patients with Sjögren's syndrome (n = 5; green, pink, purple, yellow, and blue), systemic lupus erythematosus (n = 1; navy), or pre-existing anti-SSA autoantibodies (n = 1; bright blue). **(b)** Temporal trajectories of TRIM21 NPX values for seven NMOSD patients with Sjögren's syndrome, systemic lupus erythematosus or pre-existing anti-SSA antibodies. Dashed lines indicate the time of first AQP4-IgG detection. For one patient (green), AQP4-IgG was detectable in all available samples; therefore, the timing of the first sample that was AQP4-IgG+ could not be determined. HC, healthy controls; NMOSD, neuromyelitis optica spectrum disorder.

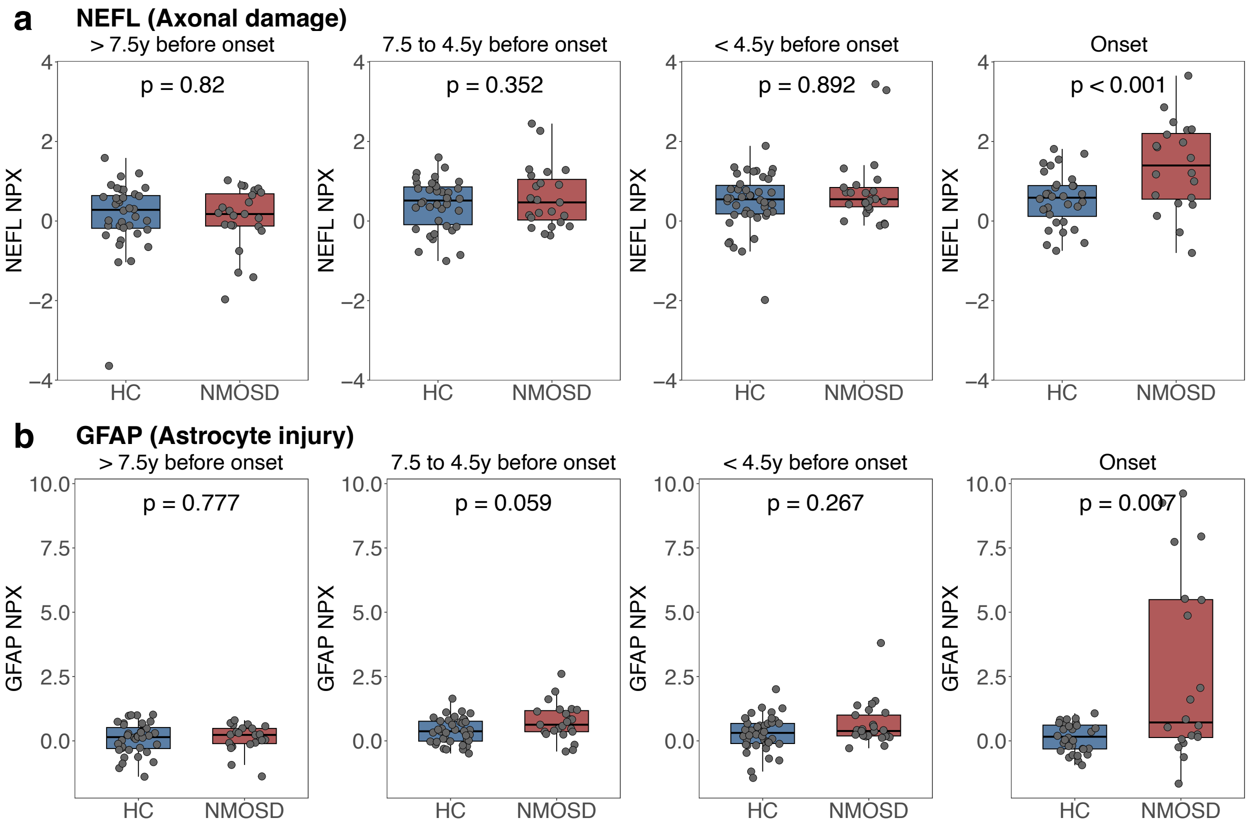


**Supplemental Figure 6. Abundance of CNS injury markers in the serum of patients with NMOSD and HC across different time intervals.** Boxplots display the median (center line), interquartile range (box), and 1.5 × the interquartile range (whiskers). Nominal p-values are derived from *limma* comparisons. GFAP, glial fibrillary acidic protein. HC, healthy controls; NEFL, neurofilament light chain; NMOSD, neuromyelitis optica spectrum disorder; NPX, normalized protein expression; y; years.

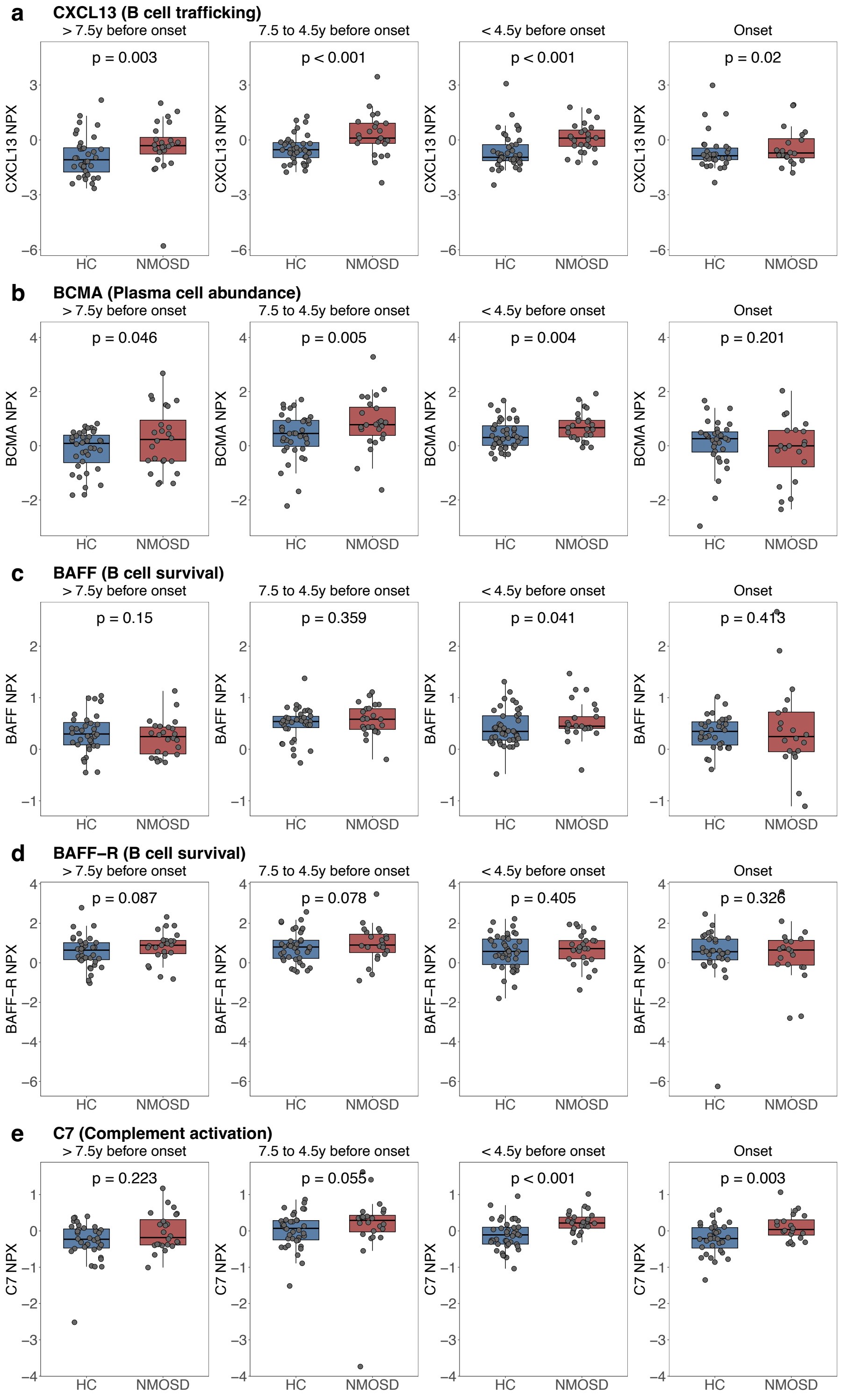


**Supplemental Figure 7. Abundance of B cell markers and complement component C7 in the serum of patients with NMOSD and HC across different time intervals.** Boxplots display the median (center line), interquartile range (box), and 1.5 × the interquartile range (whiskers). Nominal p-values are derived from *limma* comparisons. BAFF(-R), B-cell activating factor (receptor); BCMA, B-cell maturation antigen; HC, healthy controls; NMOSD, neuromyelitis optica spectrum disorder; NPX, normalized protein expression; y; years.

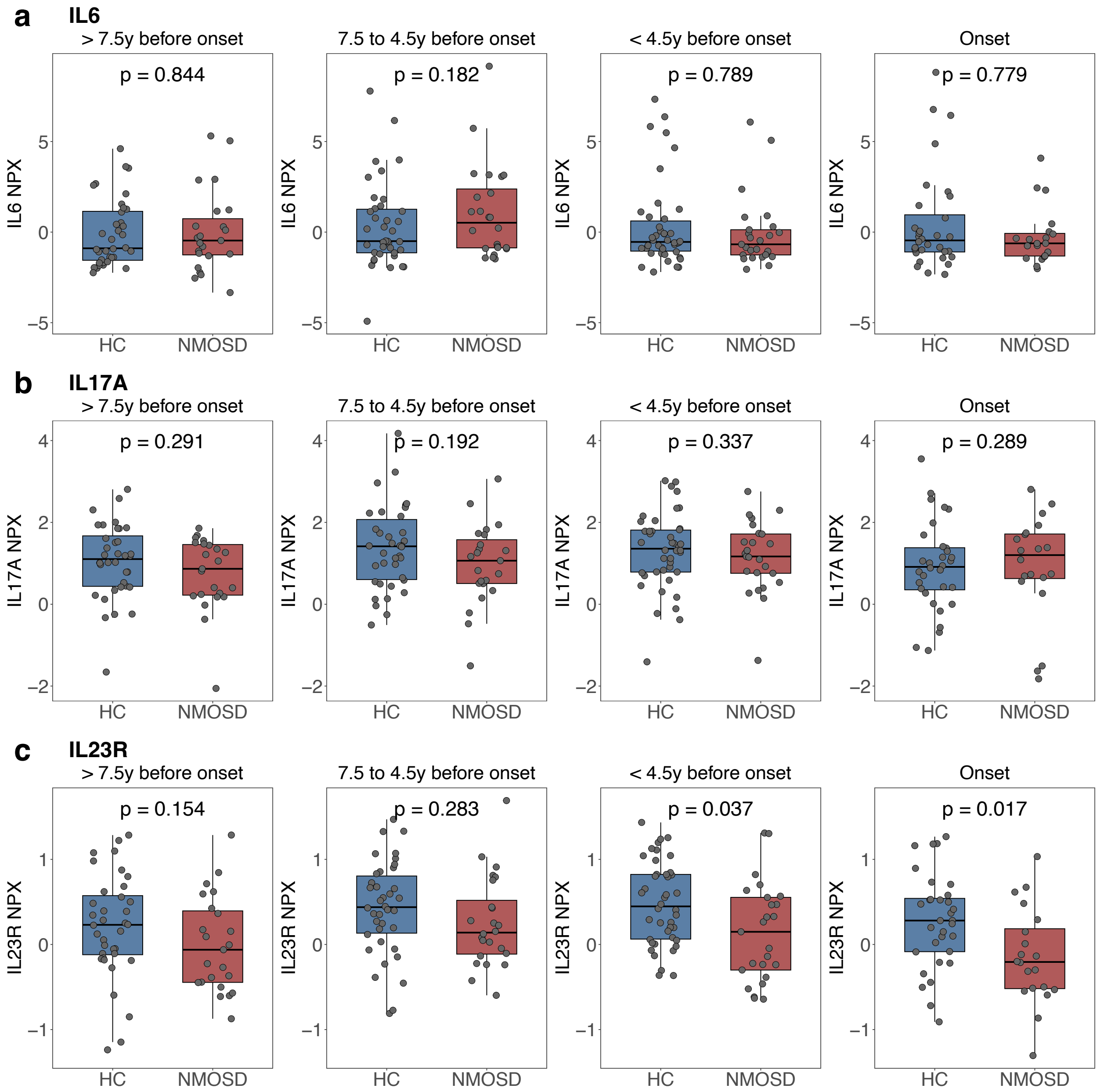


**Supplemental Figure 8. Abundance of markers related to Th17 differentiation in the serum of patients with NMOSD compared to HC.** Boxplots display the median (center line), interquartile range (box), and 1.5 × the interquartile range (whiskers). Nominal *p*-values are derived from *limma* comparisons. y; years. HC, healthy controls; NMOSD, neuromyelitis optica spectrum disorder; NPX, normalized protein expression; y; years.

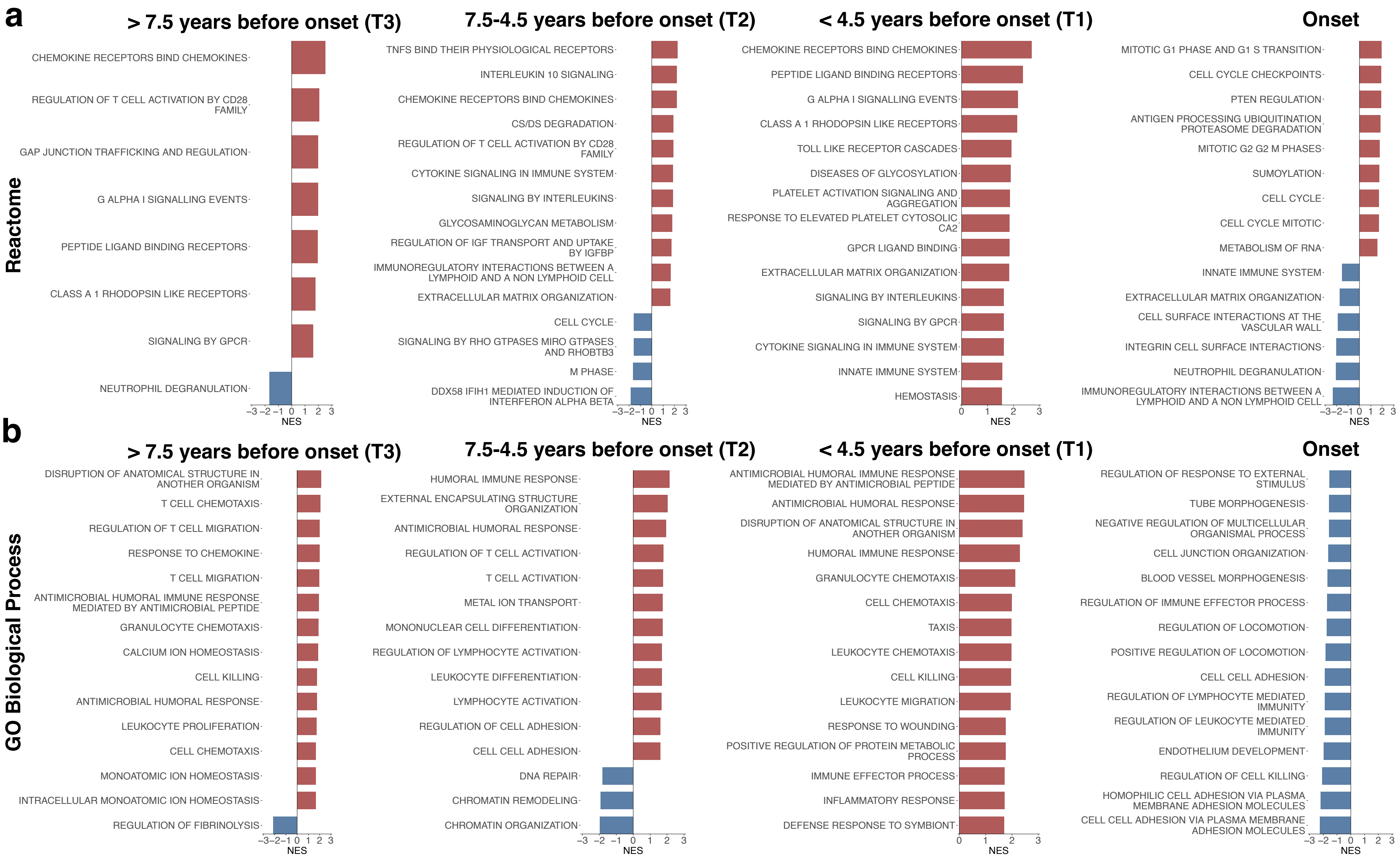


**Supplemental Figure 9: Proteome pathway analysis using the GO:BP and Reactome databases in NMOSD versus HC across time.** Gene set enrichment analysis (GSEA) was performed on ranked assays based on moderated t-statistics from *limma* analysis at onset and during preclinical stages (T1-T3). Each bar plot shows the top 15 significantly enriched pathways (FDR 10%) ranked by normalized enrichment score (NES). **(a)** Pathways from the Reactome collection. **(b)** Pathways from the Gene Ontology Biological Process (GO-BP) collection. Red and blue bars indicate pathways enriched in NMOSD and HC, respectively.

**Supplemental Table 2. Overview of five NMOSD genome-wide association study (GWAS) cohorts for *CXCL1* analysis**

| Cohort | Ancestry | N cases | N controls | Platform |
|---|---|---|---|---|
| Estrada et al. 2018[1] | EUR | 215 | 1,244 | WGS ($N_{case}$ = 86, $N_{control}$ = 460)<br>Genotyping ($N_{case}$ = 129, $N_{control}$= 784) |
| UKB | EUR | 21 | 3,416 | Genotyping |
| AoU | EUR | 62 | 6,456 | WGS |
| Yata et al. 2025[2] | EAS | 240 | 50,578 | Genotyping |
| Attfield et al. 2026[3] | Pan-ancestry | 1,573 | 1,260 | Genotyping |

EUR, European ancestry; EAS, East Asian ancestry; WGS, whole-genome sequencing; UKB, UK Biobank; AoU, All of Us.

[1] Estrada, K. *et al.* A whole-genome sequence study identifies genetic risk factors for neuromyelitis optica. *Nat Commun* **9**, 1929 (2018).

[2] Yata, T. *et al.* Contribution of germline and somatic mutations to risk of neuromyelitis optica spectrum disorder. *Cell Genomics* **5**, 100776 (2025).

[3] Attfield, K. E. *et al.* Identification of genetic risk loci associated with aquaporin 4-positive neuromyelitis optica spectrum disorder: a genome-wide association study. *The Lancet Neurology* **25**, 482–491 (2026).

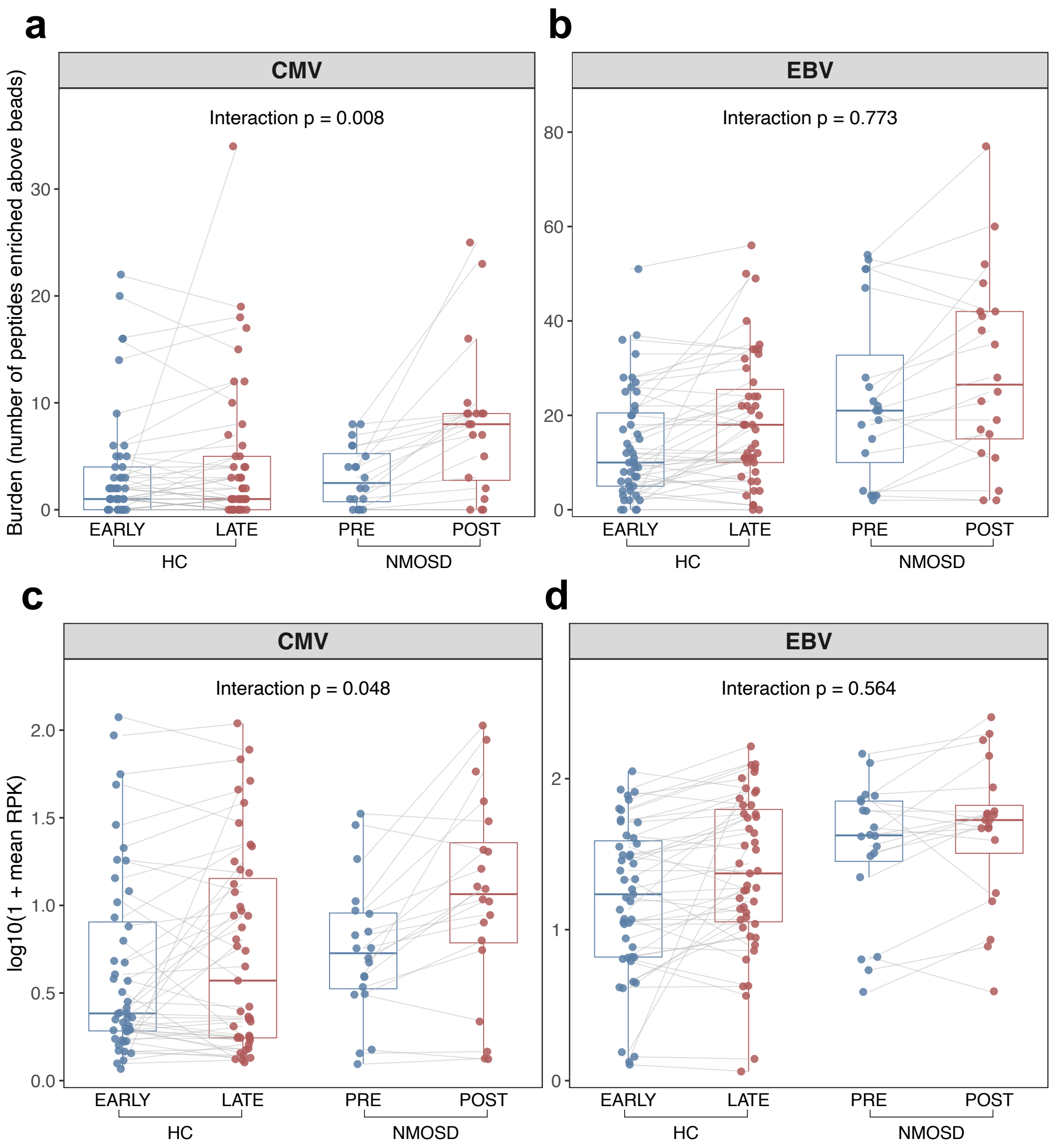


**Supplemental Figure 10: Temporal trajectories of total cytomegalovirus (CMV) and Epstein-Barr virus (EBV) antibody reactivity in patients with NMOSD versus HC.** Panels show total CMV and total EBV antibody reactivity across 47 HC with paired early/late samples and 20 NMOSD patients with paired samples collected pre- and post-AQP4-IgG seroconversion. (**a, b**) Total reactivity was defined as the number of peptides enriched above beads, across the CMV and EBV species, respectively. (**c, d**) Mean peptide signal, shown as log10(1 + mean RPK), for total CMV and total EBV reactivity, respectively. Blue and red points indicate the first and second sample per individual, respectively. Grey lines connect paired samples from the same individual. Boxplots display the median (center line), interquartile range (box), and 1.5 × the interquartile range (whiskers). Interaction p-values are derived from linear mixed-effect models and indicate whether the within-participant change over time differed between HC and NMOSD. HC, healthy controls; NMOSD, neuromyelitis optica spectrum disorder; RPK, reads per kilobase.

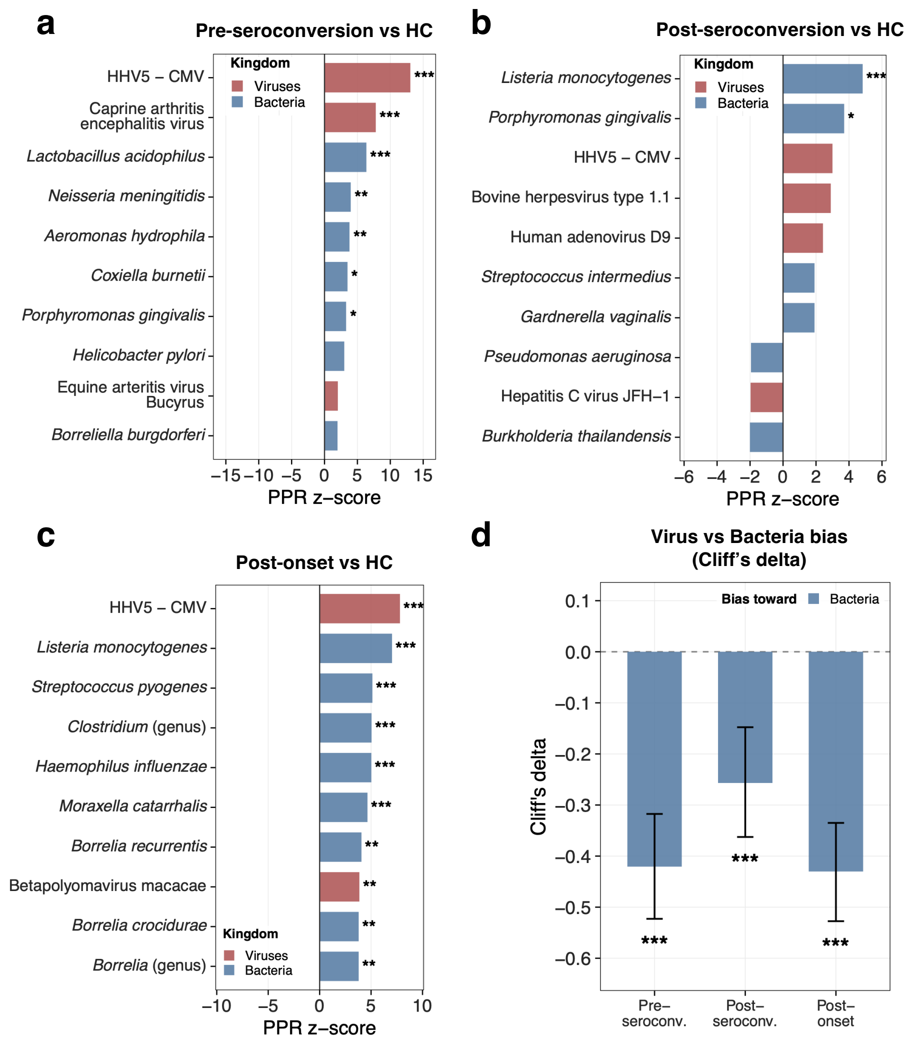


**Supplemental Figure 11: SPOKE network propagation analysis linking differentially expressed serum proteins in NMOSD to microbial organisms.** (**a-c**) Top 10 organisms ranked by z-score showing SPOKE results based on DEPs (FDR 5%) from *limma* comparisons of **(a)** pre-seroconversion preclinical NMOSD samples versus HC, **(b)** post-seroconversion preclinical NMOSD samples versus HC, and **(c)** post-onset NMOSD samples versus HC. (**d**) Cliff's δ trajectory, derived from Personalized PageRank analysis, across disease stages in NMOSD: Pre-seroconversion: δ = −0.42, 95% CI [−0.52, −0.32], $p < 0.001$. Post-seroconversion: δ = −0.26, 95% CI [−0.36, −0.15], $p < 0.001$. Post-onset: δ = −0.43, 95% CI [−0.53, −0.34], $p < 0.001$. Negative values indicate higher PPR $z$-scores for bacteria than for viruses; error bars span the 95% bootstrap CI; asterisks (* < 0.05, ** < 0.01, *** < 0.001) show organism-level Benjamini-Hochberg FDR in (**a–c**), permutation $p$-values in (**d**). Bars are colored by superkingdom (red = Viruses, blue = Bacteria); strain-level entries were aggregated to species (maximum z-score retained). CMV, cytomegalovirus; HC, healthy controls; NMOSD, neuromyelitis optica spectrum disorder; PPR, Personalized PageRank; SPOKE, Scalable Precision Medicine Open Knowledge Engine.

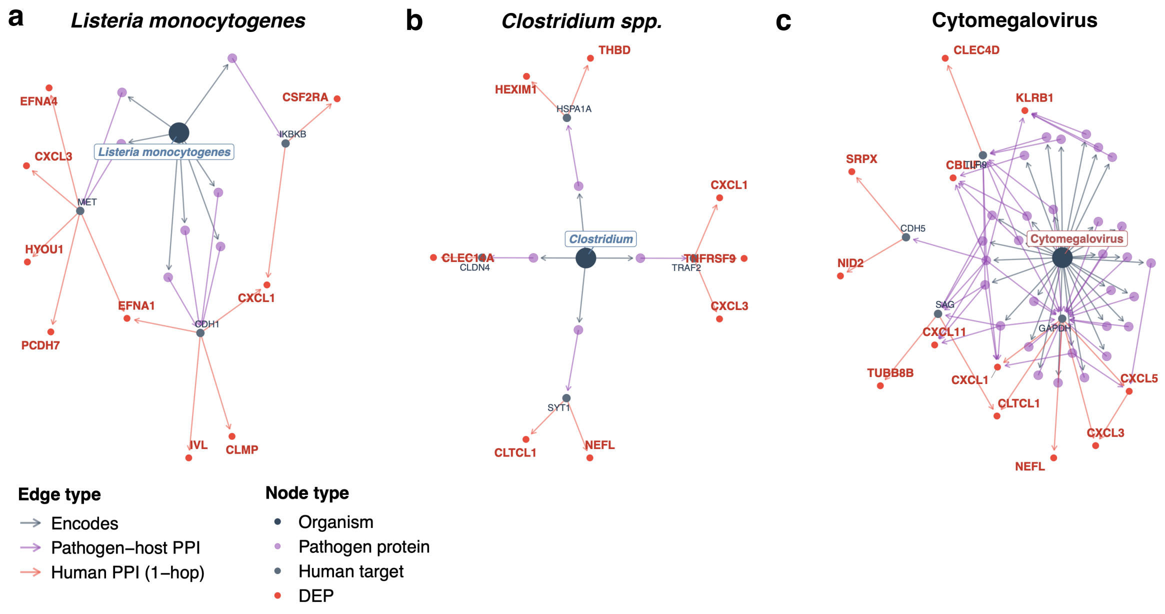


**Supplemental Figure 12: Pathogen-to-DEP metapath networks during the post-onset phase of NMOSD.** Metapaths for **(a)** *Listeria monocytogenes* (7 path proteins, 3 hubs, 9 DEPs, z = 7.04), **(b)** *Clostridium spp.* (4 path proteins, 4 hubs, 8 DEPs, z = 5.05), and **(c)** CMV (30 path proteins, 9 hubs, 12 DEPs, z = 7.81), are based on differentially expressed proteins (DEP) from *limma* group comparisons of post-onset NMOSD samples versus HC.

**Supplemental Table 3. Classifier performance in distinguishing AQP4-IgG+ NMOSD from HC or suspected seronegative cases at different timepoints.**

| Timepoint | Comparison | No. of samples | AUC | 95% CI |
|---|---|---|---|---|
| **Classifier without GFAP** | | | | |
| Onset | AQP4-IgG+ NMOSD vs. HC | 20/32 | 0.95 | 0.92-0.96* |
| T1 (< 4.5y before onset) | AQP4-IgG+ NMOSD vs. HC | 25/42 | 0.93 | 0.86-0.99 |
| T2 (4.5 to 7.5y before onset) | AQP4-IgG+ NMOSD vs. HC | 24/38 | 0.82 | 0.71-0.93 |
| T3 (> 7.5y before onset) | AQP4-IgG+ NMOSD vs. HC | 23/36 | 0.90 | 0.82-0.98 |
| Onset | AQP4-IgG+ NMOSD vs. suspected seronegative cases | 20/10 | 0.83 | 0.66-1.00 |
| **Classifier with GFAP** | | | | |
| Onset | AQP4-IgG+ NMOSD vs. HC | 20/32 | 0.95 | 0.92-0.96* |
| T1 (< 4.5y before onset) | AQP4-IgG+ NMOSD vs. HC | 25/42 | 0.93 | 0.86-0.99 |
| T2 (4.5 to 7.5y before onset) | AQP4-IgG+ NMOSD vs. HC | 24/38 | 0.82 | 0.71-0.93 |
| T3 (> 7.5y before onset) | AQP4-IgG+ NMOSD vs. HC | 23/36 | 0.90 | 0.82-0.98 |
| Onset | AQP4-IgG+ NMOSD vs. suspected seronegative cases | 20/10 | 0.83 | 0.66-1.00 |

* 95% confidence intervals (95% CI) are the 2.5th-97.5th percentiles of the fold-averaged AUC across 100 repeated 5-fold cross-validation runs; for all other timepoints, DeLong 95 % CIs are reported. AUC, area under the curve; CI, confidence interval; GFAP, glial fibrillary acidic protein; HC, healthy controls; NMOSD, neuromyelitis optica spectrum disorder;

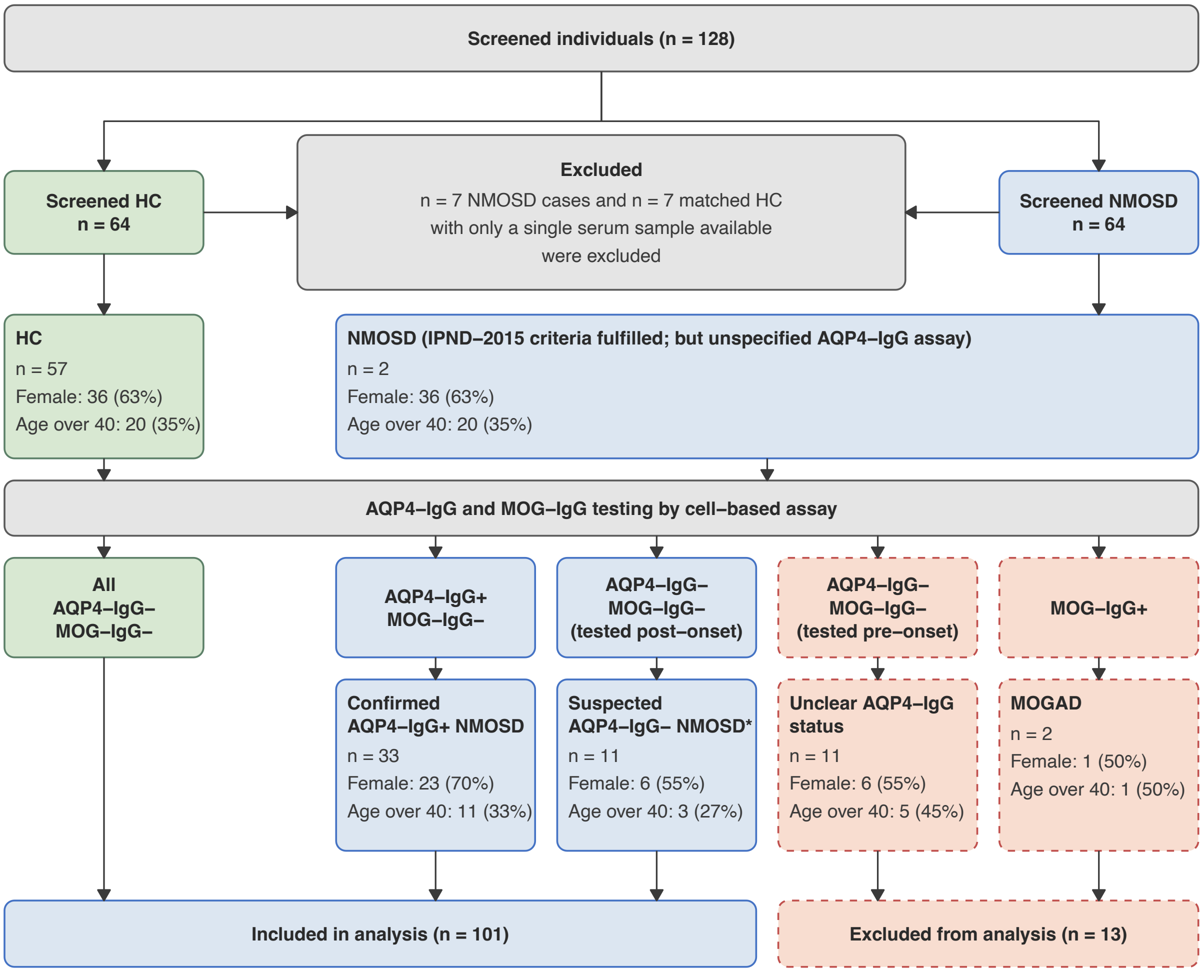


**Supplemental Figure 13. Overview of screened and excluded individuals from the U.S. Department of Defense Serum Repository.** *4 of 11 patients fulfilled the IPND-2015 criteria for seronegative NMOSD after testing by cell-based assay, the remaining 7 patients had initially tested AQP4-IgG+ by ELISA or an unspecified assay; however, this result could not be confirmed by cell-based assay on the available samples. HC, healthy controls; NMOSD, neuromyelitis optica spectrum disorder;

**Supplemental Table 4. Demographics and clinical characteristics of DoDSR cohort**

| Variable | AQP4-IgG+ NMOSD (n=33) | Suspected seronegative disease (n=11) | P-value |
|---|---|---|---|
| **Biological sex, n (%)** | | | 0.468 |
| Female | 23 (70) | 6 (55) | |
| Male | 10 (30) | 5 (45) | |
| **Age at onset, n (%)** | | | 1.000 |
| < 40 years | 22 (67) | 8 (73) | |
| ≥ 40 years | 11 (33) | 3 (27) | |
| **Presenting symptom location, n (%)*** | | | |
| Optic nerve | 12 (36) | 7 (64) | 0.164 |
| Spinal cord | 19 (58) | 5 (45) | 0.509 |
| Brain (supra- and infratentorial) | 4 (12) | 1 (9) | 1.000 |
| Area postrema | 4 (12) | 1 (9) | 1.000 |
| Unknown | 1 (3) | 0 | 1.000 |
| **Total no. of samples** | 102 | 37 | |
| Pre-onset | 82 | 27 | |
| Post-onset | 20 | 10 | |

Displayed are patient numbers with percentages. * A subset of patients had more than one symptom location at onset. P-values are derived from Fisher's Exact test.

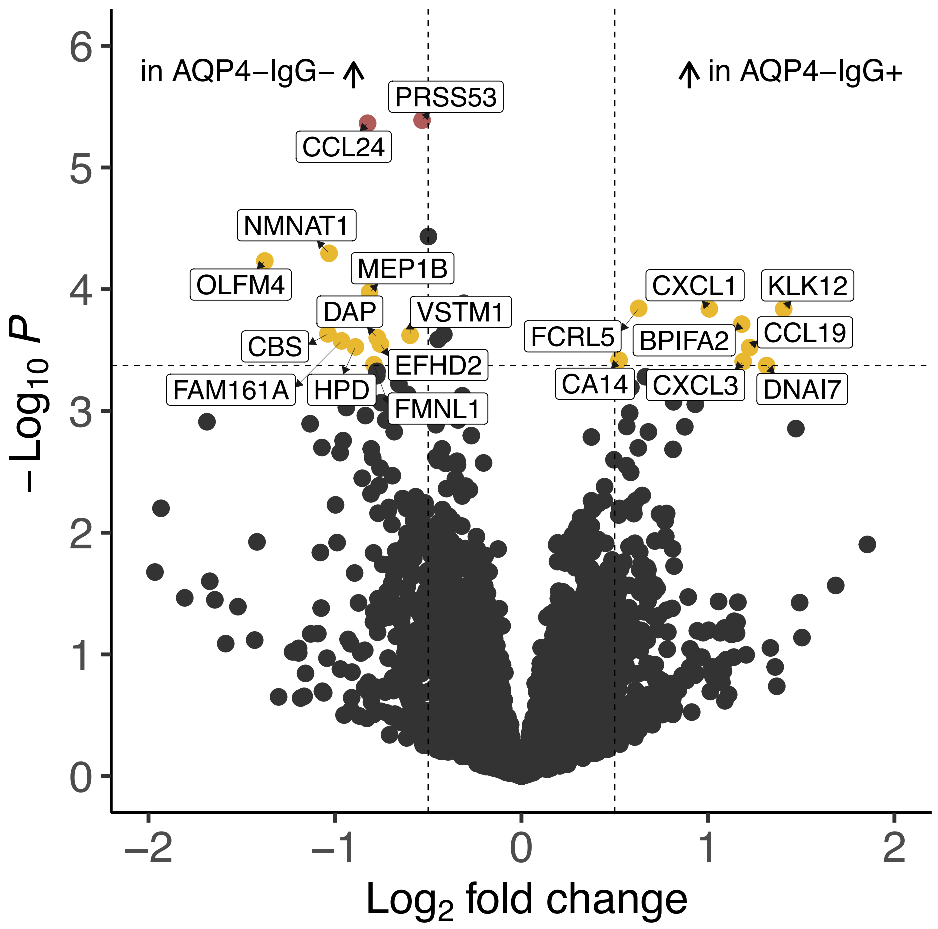


**Supplemental Figure 14: Differentially expressed serum proteins (DEP) between patients with AQP4-IgG+ NMOSD and suspected seronegative disease.** Volcano plot showing DEPs between cases of AQP4-IgG+ NMOSD (n = 102) and suspected seronegative disease (n = 37), combining samples from all time intervals (pre- and post-onset). Proteins increased in suspected seronegative NMOSD are shown on the left, proteins increased in AQP4-IgG+ NMOSD on the right. Significant DEPs at FDR 5% with a log2 fold change ($log_2FC$) > 0.5 are highlighted in red and labeled; significant DEPs at FDR 10% with a $log_2FC$ > 0.5 are highlighted in yellow and labeled. Vertical dashed lines indicate a $log_2FC$ threshold of > 0.5, horizontal dashed lines indicate an FDR threshold of 10%.

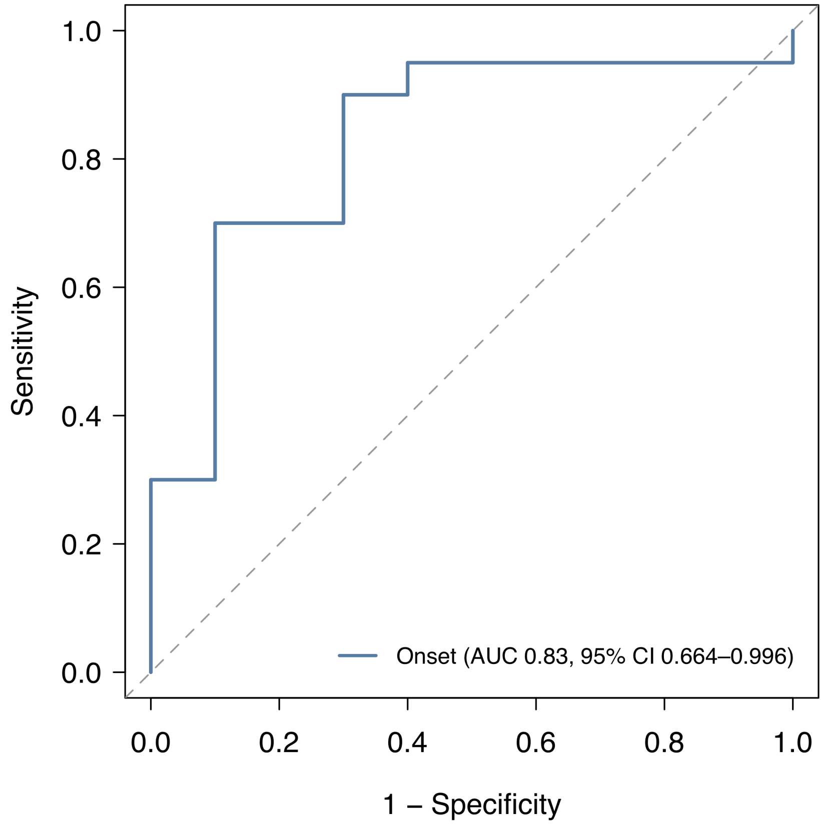


**Supplemental Figure 15. Machine-learning classifier distinguishing cases of AQP4-IgG+ NMOSD from suspected seronegative disease.** Receiver operating characteristic (ROC) curve for a ridge logistic regression model trained on post-onset samples from patients with AQP4-IgG+ NMOSD (n = 20) and healthy controls (HC; n = 32). The trained model was subsequently applied to distinguish AQP4-IgG+ NMOSD cases (n = 20) from cases of suspected seronegative disease (n = 10). 95% confidence intervals (95% CI) were calculated with the DeLong method. The dashed diagonal line indicates chance-level performance.

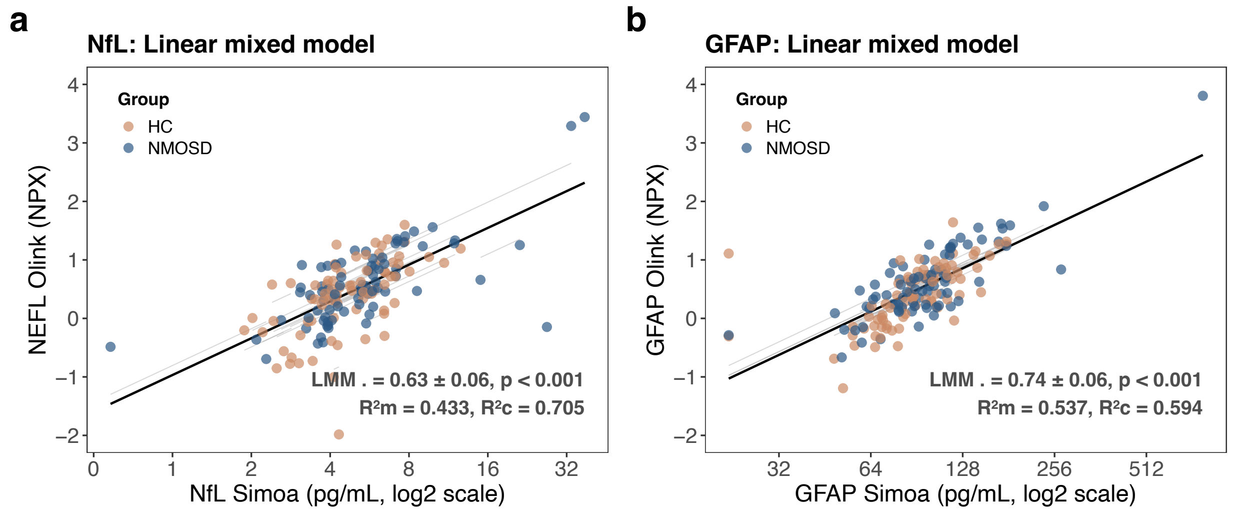


**Supplemental Figure 16. Comparison of Simoa and Olink measurements using linear mixed models (LMM).** Scatter plots show the relationship between Olink normalized protein expression (NPX) values (y-axis) and corresponding Simoa measurements (x-axis) for **(a)** neurofilament light chain (NEFL/NfL) and **(b)** glial fibrillary acidic protein (GFAP). Each point represents an individual sample, colored by group. Solid lines indicate the fitted linear mixed model (LMM). Reported statistics include the fixed-effect slope ($\beta \pm$ standard error), marginal $R^2$ ($R^2m$), conditional $R^2$ ($R^2c$), and corresponding p-values. $R^2m$ represents the variance explained by fixed effects only, whereas $R^2c$ reflects the variance explained by both fixed and random effects. HC, healthy controls; NMOSD, neuromyelitis optica spectrum disorder;
